# Depth-partitioned oxidation of atmospheric hydrogen, carbon monoxide and methane in a glacier forefield soil

**DOI:** 10.64898/2026.11.18.767582

**Authors:** Josephine Herbert, Bledina Dede, Lorenz Koschitz, Julian Sarbach, Martin H. Schroth, Marie C. Schoelmerich

## Abstract

Glacier forefields are model systems for pioneer ecosystem assembly under extreme oligotrophy. Their microbial communities carry widespread genomic potential to oxidize atmospheric trace gases, yet whether this potential is transcribed *in situ*, and by which lineages, remains largely untested. Here we combined depth-resolved profiling of hydrogen (H_2_), carbon monoxide (CO) and methane (CH_4_) at ten locations in the Wildstrubel glacier forefield, Swiss Alps, with mesocosm experiments, long-read metagenomics, and genome-resolved metatranscriptomics. Soils were consistent net sinks for atmospheric H_2_ and CH_4_, with fluxes ∼2-fold and ∼6-fold higher, respectively, in older than in younger terrain. H_2_ uptake was concentrated in the upper 20 cm, whereas CH_4_ uptake extended deeper. In the field, CO uptake was masked by apparent subsurface production, but in mesocosms lacking this source both CO and H_2_ were drawn below atmospheric concentrations. Although 57% of metagenome-assembled genomes encoded trace-gas oxidation enzymes, topsoil transcription was dominated by Group 1h [NiFe]-hydrogenases from a few Actinomycetota populations, in which the CO dehydrogenase *coxL* was transcribed 20- to 35-fold less. The second most transcribed marker was *smoX,* most closely related to C_2_-C_6_ alkane monooxygenases, suggesting the use of short-chain alkanes of possible subsurface origin. Mobile genetic elements were widely transcribed, matching host transcription, but a 104-kb plasmid-borne *coxMSL* operon was largely silent. Acetylene perturbation induced no detectable switching between trace-gas pathways. Together, these data suggest that trace-gas metabolism is partitioned across soil depth: atmospheric H_2_ scavenging dominates topsoil transcription, whereas methanotrophy and CO production occur deeper. Net surface fluxes, particularly for CO, therefore, underestimate gross trace-gas turnover.

## Introduction

Glacier forefields serve as model environments for studying primary succession and ecosystem assembly [1]. Rapid glacier retreat continuously exposes terrain to severe environmental stress, including extreme carbon and nitrogen scarcity, intense UV radiation, and broad thermal fluctuations. As soil exposure age scales predictably with distance from the ice edge, forefields form natural chronosequences ideal for tracking community establishment through time [2]. Furthermore, deglaciated terrain is rapidly expanding, projected to expose 150,000-340,000 km^2^ of new land by 2100 [3]. As soils remove 70-80% of atmospheric H_2_ [4] and are the main biological sink for atmospheric CH_4_ [5], understanding trace-gas dynamics in these emerging surfaces carries broad implications for future climate budgets.

The harsh conditions of a recently exposed forefield impose a strong selective pressure on colonization. Metagenomic and biogeochemical studies indicate that the genetic capacity for aerobic oxidation of high-energy atmospheric trace gases including molecular hydrogen (H_2_), carbon monoxide (CO) and methane (CH_4_), known as ‘aerotrophy’, is widespread in such environments, leading to the proposal that this metabolism sustains initial colonization. A substantial body of genomic and biochemical work has identified the high-affinity machineries responsible, namely Group 1 and 2a [NiFe]-hydrogenases, molybdenum-dependent CO dehydrogenases (Cox) and high-affinity particulate methane monooxygenases (pMMO) [6,7]. Trace-gas oxidation has now been documented, among others, in Antarctic deserts [8,9], arid soils [10], cave ecosystems [11] and glacial forefields [12]. Opportunistic trace-gas scavengers, primarily belonging to pioneer phyla such as Actinomycetota and Chloroflexota, establish early soil communities, generating metabolic water and organic matter that facilitate the subsequent arrival of specialized lineages [8,12]. These pioneer communities also rely on specialized chemolithoautotrophs within Pseudomonadota, which are enriched within the first years after deglaciation [13], to derive energy from mineral-bound lithic substrates such as sulfide and thiosulfate, while primary organic carbon is synthesized through ribulose-1,5-bisphosphate carboxylase/oxygenase (RuBisCO) -mediated chemoautotrophy [12]. As organic matter accumulates along the chronosequence, community composition shifts toward generalists and late-successional specialists, including methanotrophic bacteria and ammonia-oxidizing archaea like *Nitrososphaeraceae* [14,15]. Furthermore, mobile genetic elements (MGEs) play an increasingly recognized role in shaping these communities by disseminating genes of central and accessory metabolic pathways [16].

A defining feature of microbial colonization in these environments is thought to be metabolic flexibility, specifically, the ability of chemotrophic aerobes to switch between, or simultaneously co-oxidize different atmospheric trace gases (H_2_, CO and CH_4_). In laboratory settings, atmospheric methane-oxidizing bacteria (atmMOB) like *Methylocapsa gorgona* demonstrate this ability by dynamically tuning enzyme inventories, downregulating pMMO and energy-intensive biosynthesis under low-CH_4_ conditions whilst upregulating alternative trace gas machinery (i.e. [NiFe]-hydrogenase and CO dehydrogenase) to meet energetic needs [17,18]. However, while the genomic potential for multi-substrate scavenging is well-documented across harsh ecosystems [8,11,12], whether natural pioneer communities actually deploy this flexibility *in situ* remains unconfirmed. A critical gap persists between genomic capacity and environmental execution: Is active transcription distributed across multiple substrates and lineages, or concentrated in a specialized few? And does vertical spatial partitioning cause net surface fluxes to obscure deeper trace-gas dynamics? Answering these questions requires genome-resolved metatranscriptomics, which have not yet been applied to a glacier forefield.

We addressed this gap at the Wildstrubel glacier forefield in the Valais region of the Swiss Alps, a well-characterized calcareous system on the Wildhorn Nappe [14,19–21]. Wildstrubel differs from other aerotrophy study sites in that its forefield soils contain a substantial reservoir of thermogenic gas, dominated by CH_4_ (3881 ± 13 tonnes), but also containing ethane and propane, far exceeding other previously studied Swiss glacier forefields (e.g., Griessfirn forefield: 200 ± 70 tonnes) [20,21]. Where this CH_4_ is liberated by physical or chemical weathering or other disturbance, pioneer microorganisms could encounter it in addition to atmospheric trace gases. Wildstrubel therefore offers an ideal natural site for studying atmospheric trace gas oxidation and for assessing whether geologically derived reduced gases, most notably CH_4_, and short-chain alkanes, shape this activity across soil depth.

Here, we combined depth-resolved H_2_, CO and CH_4_ profiling, mesocosm and acetylene inhibition experiments, long-read metagenomics and *in situ* metatranscriptomics to determine whether, where and by whom trace gas oxidation potential is expressed. We show that trace gas metabolism is partitioned across soil depth: atmospheric H_2_ oxidation by few Actinomycetota populations dominates topsoil transcription, whereas CH_4_ oxidation and CO production occur deeper. Transcription of short-chain alkane oxidation genes further points to reduced carbon sources of possible subsurface origin.

## Materials and Methods

### Field site and sampling campaigns

Field work was performed in the Wildstrubel glacier forefield (Canton of Valais, Switzerland), a calcareous proglacial system described previously [19–21]. Two campaigns were conducted: the first conducted in 2024 is described in Supplementary Methods S1. During the final field campaign in 2024, soil was collected adjacent to location B for mesocosm experiments as described below. In early August 2025, an *in situ* acetylene inhibition experiment was conducted at site B. Soil samples (n=3) for nucleic acid extraction were collected at 5 cm depths immediately prior to treatment (sites A and B) and 24 hours post-treatment (site B only). All samples were sieved through a 1.6 mm mesh, sealed in Whirl-Pak bags, flash-frozen, transported in a dry shipper, and stored at −80 °C until processing.

### Soil gas and water-content profiles

Soil gas profiles were measured at 10 cm depth intervals through perforated access tubes using the sampling system of Nauer *et al*. (2012) [19]. Samples were stored in N_2_-flushed glass vials (CH_4_) or in Ar-flushed, modified Exetainer® vials [22]. Volumetric water content was measured at each depth in triplicate with a capacitance profile probe (PR2, Delta-T Devices); and soil temperature was logged at three sites with iButton loggers (Supplementary Methods S2).

### Soil-atmosphere flux estimation

Net soil-atmosphere fluxes (*J_net_*; positive = uptake into soil) were calculated using Fick’s first law (**eq. 3**) from concentration gradients at the soil surface (z = 0). Effective diffusion coefficients (D_eff) were derived from air-filled porosity and tortuosity determined from water content using the WLR-Marshall model [23] (total porosity θ_t_ = 0.42 [22]) alongside temperature-adjusted free-air diffusion coefficients [24] from iButton data. For CH_4_ and H_2_, surface gradients were obtained by fitting a steady-state diffusion-reaction model (**eq. 1**, [25,26]) to obtain surface gradients (**eq. 2**); full profiles were fitted except for field H_2_, which was restricted to 0-30 cm due to deeper deviations. Because CO profiles did not fit **eq. 1**, surface gradients were estimated by linear regression before applying **eq. 3**. Uncertainties were propagated from model fits and porosity and concentration measurement errors (Supplementary Methods S3).

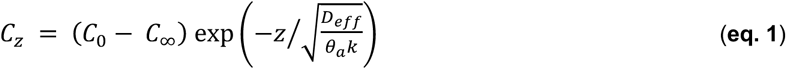

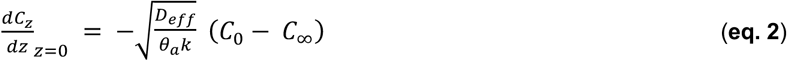

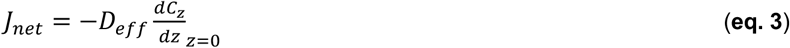

where *C_z_* is the gas concentration at depth *z* (cm); *C*_0_ is the gas concentration at the soil surface (*z* = 0); *C*_∞_ is the fitted threshold gas concentration at depth; *D*_eff_ is the effective soil diffusion coefficient (cm^2^ s^-1^); *θ_a_* is the air-filled porosity (cm^3^ cm^-3^); and *k* is the first-order consumption rate constant (s^-1^).

### Mesocosm column experiments

Approximately 20 kg of sediment was collected adjacent to site B in 10 cm depth increments to 50 cm, sieved to 8 mm and, after three days of storage at 15 °C, repacked depth-by-depth in Plexiglas columns (11 cm diameter; 80 cm height, 15 °C) to preserve the vertical stratification. Columns were fitted with gas-sampling ports every 5 cm, and time-domain-reflectometry water content sensors every 10 cm, and their headspaces were continuously flushed with compressed air (Supplementary Methods S4).

### Acetylene treatment

Acetylene (1.15 vol% in air; ALL IN GAS, Starnberg, Germany) served as inhibitor in all experiments. In the field, the acetylene mixture was delivered through ground access tubes at ∼0.5 L min⁻¹ for 2 h. Soil-gas profiles were measured before and after treatment (sites C and F in 2024; site B in 2025, with paired nucleic acid sampling). In the column, acetylene was delivered through the lowest sampling port into the basal quartz sand at 0.5 L min⁻¹ for 2 h, followed by flushing with synthetic air for 15 min. Soil for DNA-based community profiling was collected from the top 5 cm at weeks 0, 1, 3, and 10, together with gas and water-content profiles (Supplementary Methods S5).

### Apparent uptake kinetics

Apparent Michaelis-Menten parameters for H_2_ and CO consumption were estimated from serum-bottle incubations of site B soil (∼30 g, ∼10% water content, 15 °C) by following the decline of elevated headspace concentrations (∼3 or ∼6 ppm) over 5-7 h on three days, yielding consumption rates across a headspace concentration range of ∼60-7,000 ppb. Parameters were estimated by nonlinear curve fitting [27] and by weighted Lineweaver-Burk regression, and specific affinities were calculated as a⁰_s_ = V_max(app)_/K_m(app)_ [28,29]. Apparent CH_4_ uptake kinetics were derived from the mesocosm column by flushing its headspace with ∼2-100 ppm CH_4_ in hydrocarbon-free air and quantifying depth-resolved mean concentrations, fluxes and turnover rates after 18 h of equilibration (Supplementary Methods S6).

### Gas composition analysis

H_2_ and CO were quantified with a reducing-compound photometer GC (Peak Performer 1 RCP, PEAK Laboratories, CA, USA) and CH_4_ with a GC-FID (8610C, SRI Instruments Europe, Bad Honnef, Germany). Both instruments were calibrated against reference gases at the beginning of each measuring day (Supplementary Methods S7). Statistical analyses are described in Supplementary Methods S8.

### DNA and RNA Extraction

Total RNA and DNA were co-extracted from 4-5 g of soil using the RNeasy PowerSoil Total RNA Kit and the RNeasy DNA Elution Kit (Qiagen, Hilden, Germany) using a modified lysis protocol with dNTP, lysozyme and phenol/chloroform steps (Supplementary Methods S9).

### Sequencing

To obtain metagenomic data, DNA from the four mesocosm samples and three field samples underwent library preparation using the Version 14 Ligation Barcoding Kit (Oxford Nanopore Technologies (ONT), Oxford, UK; SQK-NBD114.24) and the resulting libraries were loaded to PromethION R10.4 flow cells before sequencing using the PromethION 2 Solo platform (ONT, Oxford, UK; PRO-SEQ002). Basecalling was done using the Super Accurate model (<u>dna_r10.4.1_e8.2_400bps_sup@v5.2.0</u>) using Dorado v7.9.8 through the MinKnow v25.05.12 GUI interface.

For metatranscriptomic data generation for the field samples before and after acetylene inhibition, sequencing was performed by Novogene (Novogene Germany GmbH, Münich, Germany) using the Illumina NovaSeq X Plus platform (paired-end 150 bp).

### Metagenomic data processing and MAG recovery

Raw metagenomic data obtained from ONT sequencing first underwent quality assessment and control (quality-score >15, length >100 bp) using Chopper v0.10.0b [30], and adapters were removed using Porechop v0.2.4 [31]. Assemblies were created using Flye v2.9 (--meta, --nano-hq and -- iterations 3) [32]. To assess the relative abundance of taxa from the filtered reads, Kraken2 v2.1.3 and Bracken v3.0.1 [33,34] were used against the GTDB r220 database (09/11/2024).

Metagenome-assembled genomes (MAGs) were generated as described in Fiorito *et al*., 2026 [35], dereplicated at 95% ANI using dRep v3.4.5 [36] to only retain species-level representatives, and their relative abundance estimated using CoverM v0.6.1 [37] to map the filtered reads back to the dereplicated genomes. Due to poor MAG recovery, further MAGs were generated from a co-assembly of all samples, assembled using the same methodology as above, followed by the same binning methodology and dereplication. For their taxonomic classification, GTDB-Tk v2.4.0 [38] was employed, using the GTDB r220 database (Supplementary Methods S10).

Unbinned contigs from all seven per-sample assemblies and the co-assembly were isolated by removing MAG-assigned contigs identified via DAS Tool v1.1.7 [39], excluding sequences aligning to dereplicated MAGs at ≥80% query coverage using minimap2 v2.22 [40], and dereplicating the remainder at 99% nucleotide identity (80% coverage) using MMseqs2 v14 [41] (Supplementary Methods S10).

For functional annotation, Prodigal v2.6.3 [42] was used to generate protein profiles. Metabolic marker genes were searched using a curated protein database of 59 metabolic marker genes (https://github.com/GreeningLab/GreeningLab-database; Accessed 09.10.2026). These were searched as described in Ricci *et al*., 2025 [12] with more details within Supplementary Methods S10. Hydrogenases were identified using HMMsearch (HMMer v3.4) [43] using the HydDB database (2019) [44] (Supplementary Methods S10). To confirm the phylogeny of certain marker genes, namely *smoX*, *coxL* and *pmoA/amoA*, phylogenetic trees were constructed, described further in Supplementary Methods S11.

### MGE Identification, annotation and activity analysis

MGEs were identified in the same manner as Dede *et al*., 2026 [45], described further in detail in Supplementary Methods S12. Their activity was assessed by using CoverM v0.6.1 [37] to map the metatranscriptome to the vOTU and pOTU contigs for whole activity, and then applying the TranscriptM v0.5 pipeline for resolving marker gene transcription as described below. Host linkage was determined using iPHoP v1.4.1 [46] with MAGs added to the database (June 2025 release, GTDB r220) for closer associations.

### Metatranscriptomic data processing and analysis

Metatranscriptomic data was processed by first undergoing quality control using FastQC v0.23.4, followed by the TranscriptM v0.5 [47] pipeline as performed in Dede *et al.*, 2026 [45]. Transcripts per million (TPM) values were log-transformed as log_10_(TPM+1) where zero values occur.

## Results

### The Wildstrubel glacier forefield is a consistent H2 and CH4 sink that strengthens with soil age

To characterize depth-resolved gas dynamics and quantify trace gas fluxes across the Wildstrubel forefield, soil gas and moisture profiles were sampled across ten locations spanning two successional stages: young (∼20 years post-deglaciation) and older (∼50 years post-deglaciation) terrain (**Figure 1A**). Molecular H_2_ mixing ratios consistently decreased with soil depth, establishing the Wildstrubel glacier forefield as a robust and continuous H_2_ sink. The vast majority of H_2_ consumption was concentrated within the upper 10-20 cm of the soil profile, below which concentrations stabilized or exhibited minor increases (**Figure 1B**), matching depth-stratified oxidation profiles observed by us and others in other polar and alpine soils [8,12,19]. The magnitude of this biological sink was strongly coupled to the successional stage of the forefield; the older terrain exhibited a mean H_2_ consumption flux of 275 µmol m^-2^ d^-1^, nearly double that of the younger terrain (155 µmol m^-2^ d^-1^) (**Figure 2A**). Trace-gas oxidation likewise increased with soil age in an Antarctic foreland, whereas *ex situ* H_2_ oxidation rates were invariant across the Swiss Griessfirn chronosequence [12]. Our absolute H_2_ fluxes were lower than the 815 µmol m^-2^ d^-1^ reported at Griessfirn, likely reflecting local thermal regimes and soil matrix properties known to influence trace-gas uptake [48].

**Figure 1.**
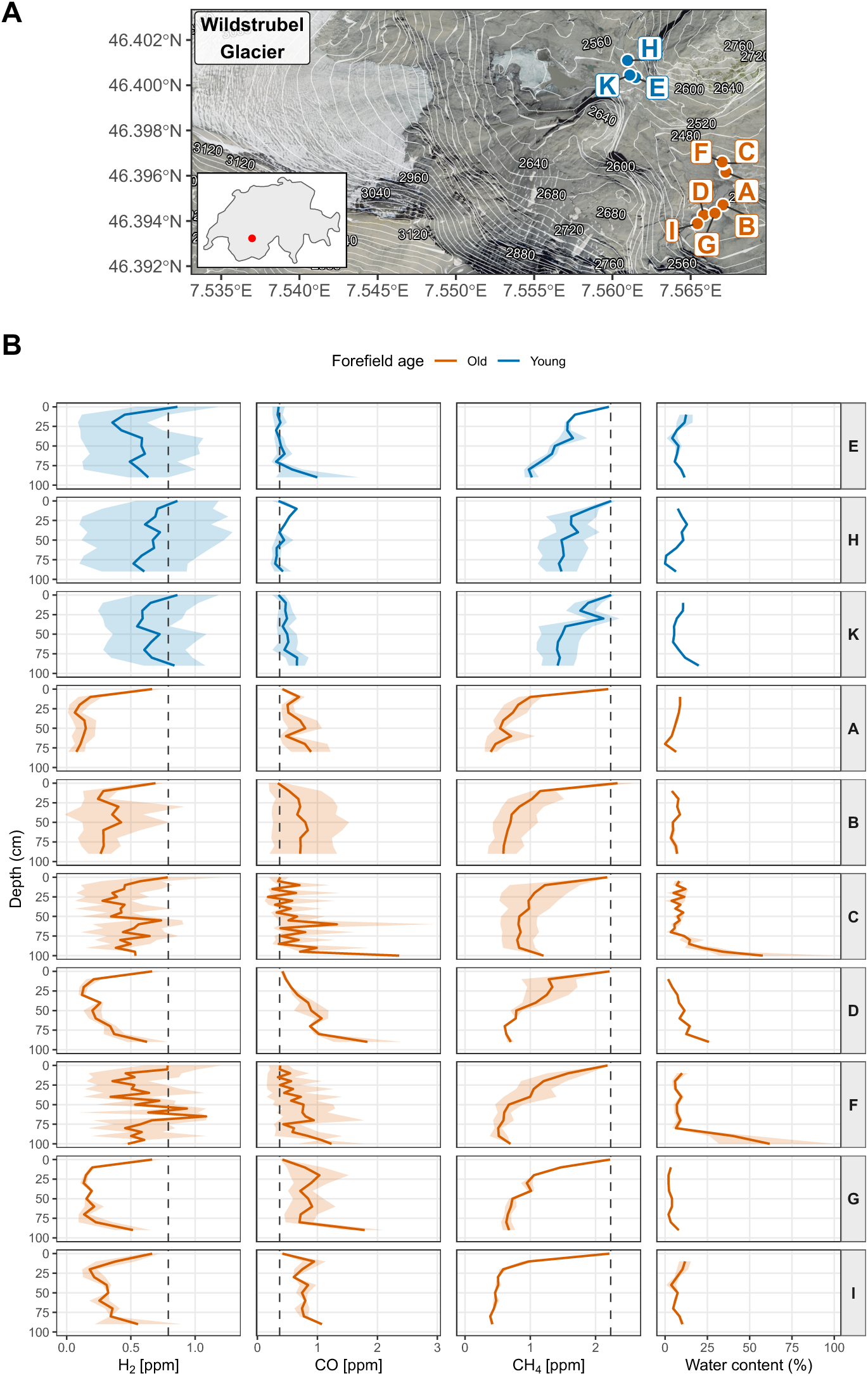
Trace-gas depth profiles across the Wildstrubel glacier forefield chronosequence. **(A)** The Wildstrubel forefield, Switzerland (Bernese/Valaisian Alps; location within Switzerland shown in inset map as red dot), shown on a SWISSIMAGE aerial orthophoto (swisstopo) with 20 m elevation contours (white, derived from the swissalti3d digital elevation model). Circles mark the ten sampling sites, colored by deglaciation history: a younger section deglaciated approximately 20 years before sampling (E, H, K; blue) and an older section deglaciated approximately 50 years before (A-D, F, G, I; orange). Coordinates are shown as CH1903+/LV95 (EPSG:2056). Base data: swisstopo. **(B)** Depth profiles of H2, CO and CH4 (ppm) and volumetric water content (%), grouped by soil age. Lines show the mean at each depth across all sampling campaigns at that site (n = 2 campaigns at sites A, D, E, G, H, I and K, n = 3 at B, n = 5 at C and F; 27 profiles in total for H2, CO and water content, and 21 for CH4, which has 2 per site; 26 August-3 October 2024); ribbons give the full observed range. Dashed vertical lines show the mean ambient-air mixing ratio (H2 = 0.79, CO = 0.37, CH4 = 2.23 ppm; n = 21, 21 and 16 ambient samples, respectively). Colors reflect the same as in A.

**Figure 2.**
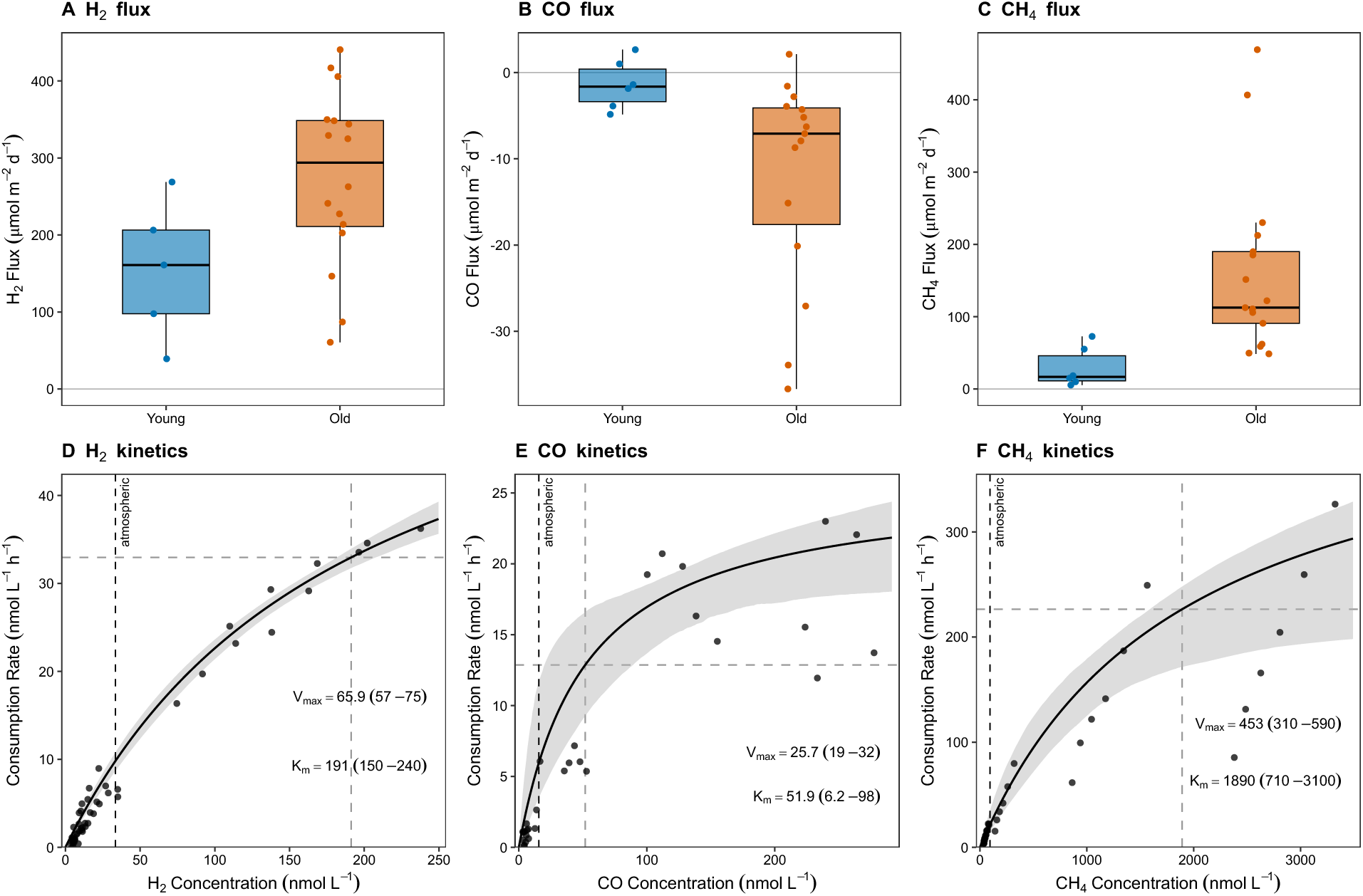
Soil-atmosphere trace-gas fluxes and apparent uptake kinetics. **(A-C)** Net uptake fluxes of H2 **(A)**, CO **(B)** and CH4 **(C)** in the older (sites A-D, F, G, I) and younger (sites E, H, K) forefield, calculated from field soil-gas profiles using Fick’s first law, with the effective diffusion coefficient derived from volumetric water content, a porosity of 0.42 and soil temperature (see Methods). Acetylene-inhibited profiles and unreliable fits were excluded (H2: older n = 16, younger n = 5; CO: older n = 15, younger n = 6; CH4: older n = 17, younger n = 6). **(D-F)** Apparent uptake kinetics of H2 **(D)**, CO **(E)** and CH4 **(F)** in forefield sediment at 15 °C, showing consumption rate per litre of incubation volume against concentration (H2 n = 51, CO n = 30, CH4 n = 36). Curves are Michaelis-Menten functions from weighted Lineweaver-Burk regressions with 95% bootstrap confidence bands; Vmax and Km (95% confidence interval) are annotated. Grey dashed lines mark Km(app) and Vmax(app). The black dashed line marks the mean ambient concentration (H2 = 33.5, CO = 15.7, CH4 = 94 nmol L⁻¹). Flux summaries and full kinetic parameters are in Tables S13 and S14; Lineweaver–Burk fits are in Supp. Figure 1.

CH_4_ displayed a similar consumption pattern, with all ten locations being net atmospheric sinks under undisturbed conditions. Concentration profiles declined exponentially from ambient mixing ratios (mean 2.23 ± 0.10 ppm (n = 16)) near the surface to minimal values at depth (**Figure 1B**), with the steepest uptake gradients again localized in the top 20 cm. Net CH_4_ consumption scaled strongly with soil age: older forefield soils yielded a median net flux of 113 µmol m^-2^ d^-1^ (n = 17), compared to 17 µmol m^-2^ d^-1^ in younger terrain (n = 6) (Wilcoxon test, p < 0.001; **Figure 2C**). Within the older terrain, geomorphology further modulated gas dynamics, with sandhill sites (sites A, B, D, G and I) displaying ∼45% higher net uptake than terrace locations (sites C and F) (median 122 vs 84 µmol m^-2^ d^-1^), in line with adjacent Swiss forefields [14,19].

In contrast, subsurface CO concentrations increased with depth in older forefield soils, rendering them a net atmospheric CO source, whereas the younger terrain maintained near-zero net flux (**Figure 2B**). Source strength in older soils varied more across sampling dates than across geographic locations, pointing to a moisture- or temperature-dependent abiotic mechanism, such as water table dynamics (prominent at sites C and F) or the potential release of soil-entrapped CO via bedrock weathering [48]. Because subsurface abiotic production and surface biological oxidation operate in opposing directions, net surface CO fluxes systematically underestimate gross biological uptake, and near-zero net fluxes in younger soils reflect process cancellation rather than inactivity, as observed in other glacial systems [11,12,48].

### H2, CO and CH4 are oxidized at sub-atmospheric concentrations in older forefield soils

To determine whether forefield microbial communities sustain trace-gas oxidation under substrate-limiting conditions and to characterize their uptake kinetics, apparent enzyme kinetics were measured in laboratory serum bottles containing soils from our focal site B. Lineweaver-Burk analysis showed nanomolar-scale uptake kinetics for both H_2_ (V_max(app)_ = 65.9 nmol L^-1^h^-1^; K_m(app)_ = 191 nmol L^-1^; a⁰_s_ = 0.345 h^-1^) and CO (V_max(app)_ = 25.7 nmol L^-1^ h^-1^; K_m(app)_ = 51.9 nmol L^-1^; a⁰_s_ = 0.496 h^-1^), with H_2_ consumed faster overall but CO scavenged more efficiently at trace concentrations (**Figure 2D, E, Supp. Figure 1**). Sub-atmospheric CH_4_ consumption at site B was likewise observed, declining from 99 (near ambient) to ∼29 nmol L^-1^ by 50 cm depth, with active topsoil (0-30 cm) kinetics of K_m(app)_ = 1,890 nmol L^-1^ and V_max(app)_ = 453 nmol L^-1^ h^-1^ (**Figure 2F, Supp. Figure 1**). All three gases were therefore oxidized under substrate-limiting, sub-ambient conditions, establishing that the microbial machinery required for atmospheric concentration scavenging was present and functional.

### Genomic potential for trace-gas oxidation is broadly distributed across the community

To investigate the community underlying these fluxes, we generated metagenomes from the topsoil (5 cm) in the older forefield (∼50 years post-deglaciation; site B), the successional stage with the highest trace-gas uptake and demonstrated sub-atmospheric oxidation. MAGs were reconstructed from the field (n=3) and mesocosm (n=4), resulting in 351 initial bins, and a further 219 bins via co-assembly, leading to 159 MAGs dereplicated at species-level (>=95% ANI; 34 high-quality (HQ) and 125 medium quality (MQ)) by MIMAG standards [49]. These consisted of 157 bacterial MAGs across 11 phyla and 2 archaeal MAGs (both Thermoproteota). The resulting community architecture, dominated by Actinomycetota, Chloroflexota and Pseudomonadota, matches established post-glacial chronosequences at intermediate successional stages (∼50 years) [12,50] (**Figure 3A**). Screening these genomes for trace-gas oxidation markers revealed that 91 of 159 MAGs (57.2%) encoded at least one Group 1 or 2a [NiFe]-hydrogenase or *coxL* (**Supp.** Figure 5), with 18 encoding markers for two or more substrates (**Figure 3B**). This capacity spanned six phyla rather than confined to a single lineage.

**Figure 3.**
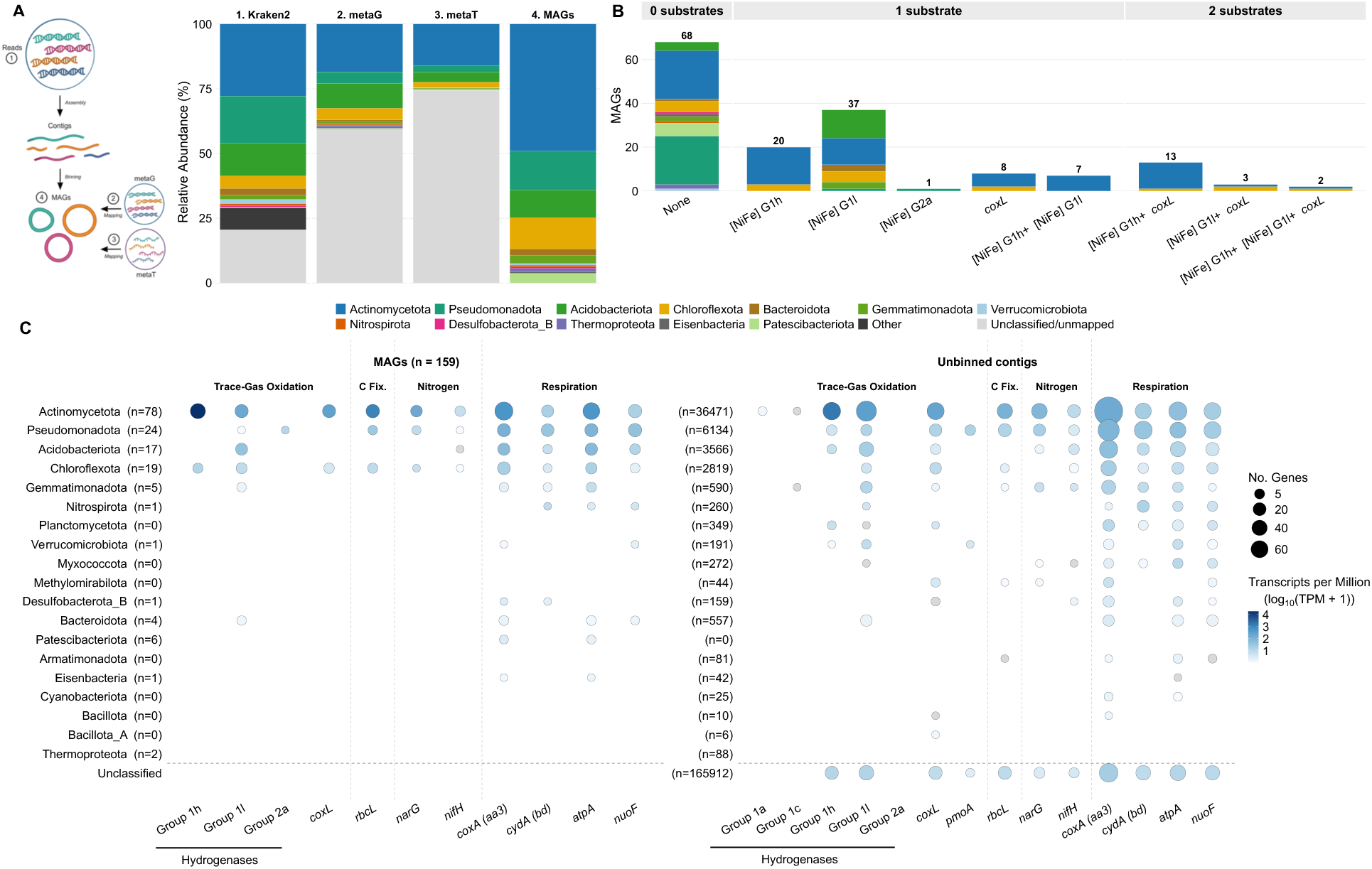
Community composition and encoded trace-gas oxidation potential. **(A)** Schematic of MAG recovery pipeline and corresponding analyses followed by stacked bar charts showing 1) the relative abundance of bacterial and archaeal phyla by read-level classification, 2) metagenomic read-mapping to MAGs, 3) metatranscriptomic read-mapping to MAGs, and 4) phyla composition of the recovered MAGs. Grey denotes reads unclassified at phylum level in the first column and reads not mapping to any MAG in the two mapping columns. Note that the first three columns are abundance-weighted whereas the fourth counts genomes. Phyla not within MAGs collapsed into ‘Other’. **(B)** Number of trace-gas markers (Group 1a, 1c, 1h, 1l, and 2a [NiFe]-hydrogenases, *coxL, pmoA*) encoded per MAG grouped by number of substrates (H2, CO, CH4), colored by phylum. **(C)** Transcription of marker genes across phyla, shown separately for MAGs and for unbinned contigs, separated by phylum with n denoting the number of MAGs or contigs identified within that phylum. Circle area gives the number of genes within each phylum; fill gives summed transcript abundance (TPM, log10), grey fill indicates a TPM of zero. Markers are grouped by function (C fix. = carbon fixation); *coxL* denotes the CO dehydrogenase large subunit; *pmoA* = particulate methane monooxygenase subunit A; *rbcL* = ribulose-1,5-bisphosphate carboxylase/oxygenase (RuBisCO); *narG* = dissimilatory nitrate reductase; *nifH* = nitrogenase; *coxA* = cytochrome bo3 oxidase; *cydA* = cytochrome bd oxidase; *atpA* = ATP synthase; *nuoF* = complex F I subunit. Transcript data derived from a single native field library (WIL2 Before; WBe15_8).

MAGs captured only part of the community. Since 59.1% of metagenomic reads did not map to these MAGs (**Figure 3A**), we also screened the unbinned fraction of contigs. Markers were recovered in comparable proportions and within similar phyla (**Figure 3C**), indicating that the MAG-based inventory reflects the wider community. The exception was *pmoA*, since only fourteen *pmoA* genes were recovered on unbinned contigs but none in the 159 MAGs (**Figure 3C, Supp. Figure 5**). Canonical pMMO-encoding methanotrophs were thus present at 5 cm but too rare to be recovered as MAGs, consistent with a methanotrophic layer below our sampling depth.

### Active transcription is narrow, H2-dominated, and concentrated in few Actinomycetota populations

To determine active gene expression, we generated metatranscriptomes from the same site B topsoil, comprising two libraries collected immediately before and 24 h after acetylene inhibition *in situ* (see Methods and below). Although lacking biological replication, these represent, to our knowledge, the first metatranscriptomes from a glacier forefield that resolve trace-gas metabolism [13]. Because both libraries capture a snapshot rather than a steady state, our analysis relies on within-genome expression ratios, which are robust to library effects.

The dominant transcriptional signal was consistent across both libraries (**Supp. Figure 3**), but here we describe only the pre-treatment library, to exclude any effect of acetylene inhibition. Among the 59 markers screened, Group 1h (G1h) [NiFe]-hydrogenases were the most highly transcribed, with four Actinomycetota MAGs of the genus CADCVQ01 (*Solirubrobacterales*, Actinomycetota) accounting for 69.6% of G1h transcripts (**Figure 4A, C; Supp. Figures 2A, 4A**). This genus is unnamed and so we propose a name within Supplementary Results S1. Some of these MAGs also transcribed RuBisCO, central for carbon fixation via the Calvin-Benson-Bassham (CBB) cycle (Supplementary Results S2). Unbinned contigs mirrored this dominance, with 142 G1h hydrogenases present on 100 contigs (Actinomycetota 66, unclassified 20, Pseudomonadota 7, Other 7), and 315 Group 1l (G1l) hydrogenases on 212 contigs (Actinomycetota 104, Acidobacteriota 32, Unclassified 31, Other 45), and G1h hydrogenases representing the top transcribed marker (**Figure 4A, D**), with a single CADCVQ01 contig accounting for 76% of G1h hydrogenase transcripts (**Supp. Figures 4D, 2B**). High-affinity [NiFe]-hydrogenase transcription was therefore concentrated in a few Actinomycetota lineages.

**Figure 4.**
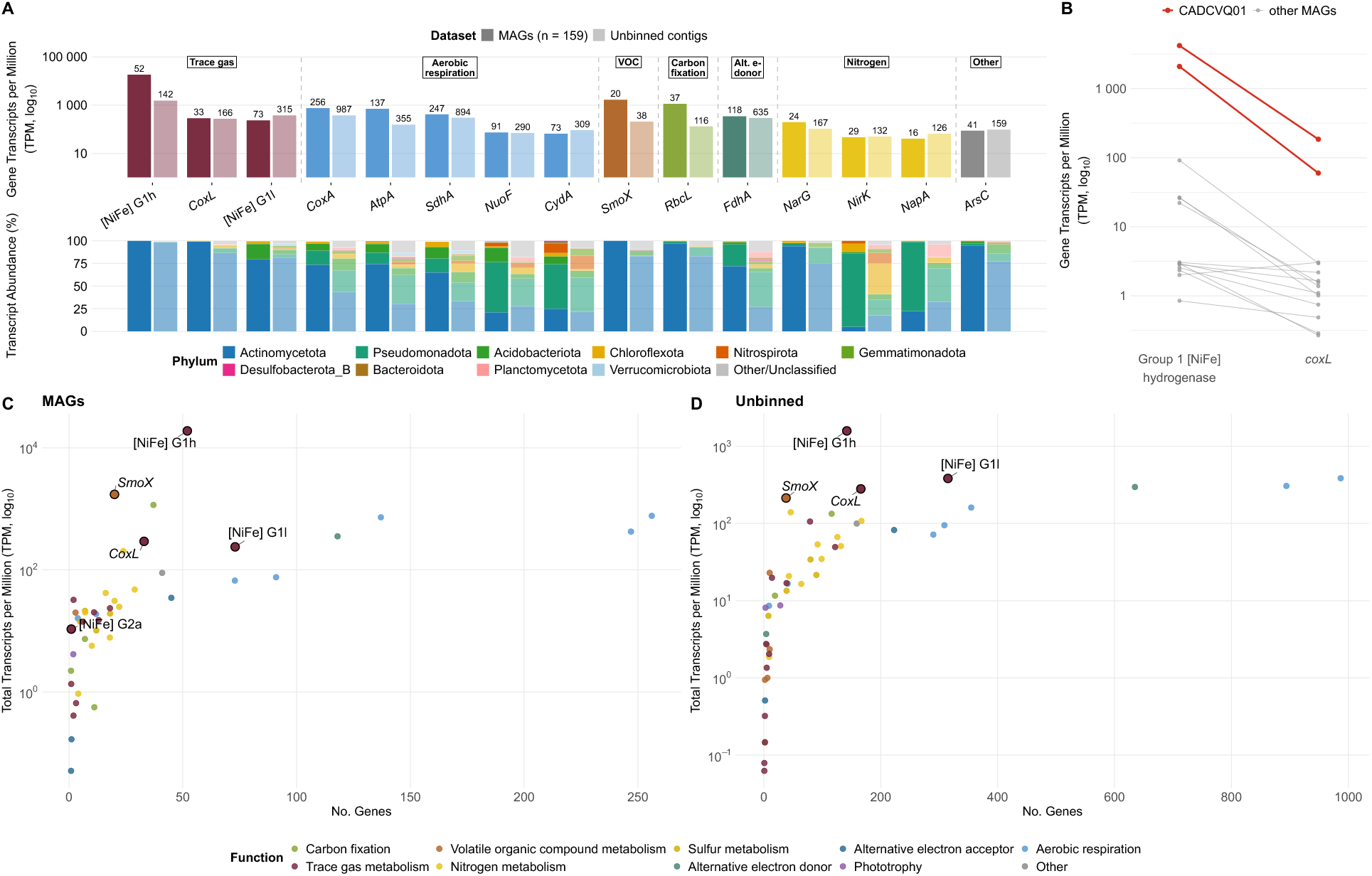
Transcription of trace-gas oxidation machinery is narrow and hydrogen-dominated. **(A)** The top 15 most transcribed functional genes in the native field library, ranked by transcript abundance (Transcripts per Million; TPM, log10) and colored by functional category. Lower strip, the taxonomic composition of transcripts assigned to each gene, expressed as a percentage of transcripts mapping to MAGs or unbinned contigs. Darker color indicates MAGs and lighter color indicates unbinned contigs. Phyla not within top 10 collapsed into ‘Other’ shown in grey, which also includes unclassified unbinned contigs not resolved to phylum, n denotes number of genes identified. **(B)** Transcript abundance of Group 1 [NiFe]-hydrogenases and *coxL* in 15 MAGs encoding both. Lines connect the two values within a single genome, so each comparison is internally controlled for library depth, extraction efficiency and genome copy number; CADCVQ01 genomes are highlighted in red. Both panels derive from a single field library (site B, 5 cm; WIL2 Before; WBe15_8). **(C)** Number of marker genes in MAGs and **(D)** unbinned contigs against their total TPM (log10), highlighted are Group 1h, 1l, and 2a [NiFe]-hydrogenases ([NiFe] G1h, [NiFe] G1l, [NiFe] G2a), large subunit of CO dehydrogenase (*coxL*) and soluble diiron monooxygenase (*smoX*). Dots colored by functional category of marker genes, matching panel A.

Among MAGs encoding both a Group 1 hydrogenase and *coxL*, hydrogenase transcription exceeded *coxL* transcription in 14 of 15 cases, and by 20- to 35-fold in the CADCVQ01 MAGs (**Figure 4B**). Community-wide *coxL* transcription followed the same pattern, i.e., detectable but consistently far lower than hydrogenase transcription levels.

### Soluble di-iron monooxygenases point to an expanded substrate range of generalist pioneers

A further, unexpected feature was the prevalence and transcription of *smoX*, encoding soluble di-iron monooxygenases (SDiMOs). We identified *smoX* in 15 MAGs (13 Actinomycetota, 2 Pseudomonadota) and 38 hits within 33 unbinned contigs. Among the 59 markers screened, *smoX* was the second most highly transcribed after G1h hydrogenases (**Figure 4A, C**). Transcription was concentrated in a single *Pseudonocardia* MAG (**Supp. Figure 4B**), but *smoX* was also encoded in JACCQS01 (*Geodermatophilales*, Actinomycetota) alongside *coxL*, *rbcL*, *fdhA*, *coxA*, and G1h/G1l hydrogenases, establishing this taxon as a metabolic generalist with the potential to contribute to dark primary production, as described for versatile Chloroflexota in alpine forefields [12]. In the unbinned fraction, *smoX* transcription was similarly concentrated in three Actinomycetota contigs, contributing 61% of its TPM (**Supp. Figure 4E**).

SDiMOs are a versatile enzyme class, capable of oxidizing substrates such as methane, short-chain alkanes, aromatics, and chlorinated pollutants [51]. Smo-type enzymes most closely related to our sequences are active on C_2_-C_6_ alkanes rather than CH_4_ (**Figure 5**), so high *smoX* expression more likely represents short-chain alkanotrophy rather than supplementary methanotrophy. This metabolic role is supported by local gas chemistry: while CH_4_ dominates the C_1_-C_3_ hydrocarbon pool (>96% in soils), short-chain alkanes are consistently present, with ethane (C_2_H_6_) contributing 0.6-2.5% and propane (C_3_H_8_) up to 0.8% of soil gases, alongside select subsurface rocks containing up to 58.6% C_2_H_6_ [21]. Transcription of a putative C_2_-C_6_ monooxygenase in soils containing entrapped alkanes suggests a previously overlooked subsurface energy source complementing atmospheric trace-gas scavenging in early forefield succession.

**Figure 5.**
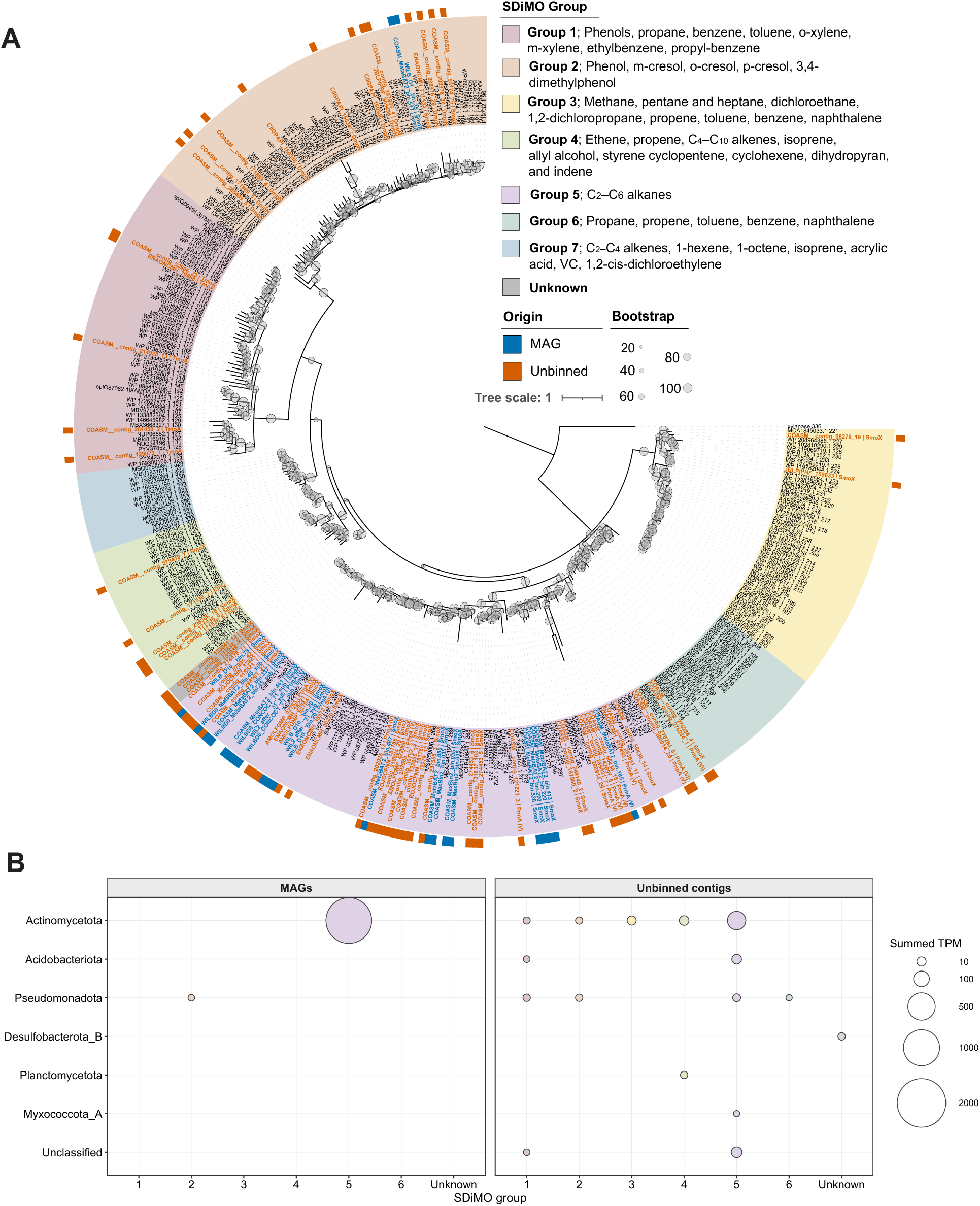
Phylogeny of recovered soluble diiron monooxygenases places the highly transcribed *smoX* genes among short-chain alkane monooxygenases. **(A)** Phylogenetic tree of all genes recovered as soluble diiron monooxygenases (SDiMOs), including *smoX*, from MAGs (n=159) and unbinned contigs and their phylogenetic grouping based on Yang *et al*., 2025. Highlighted in blue are proteins from MAGs and in orange from unbinned contigs. Reported substrates of each SDiMO group are: Group 1; phenols, propane, benzene, toluene, o-xylene, m-xylene, ethylbenzene and propylbenzene; Group 2; phenol (Shingler *et al*., 1989) [52], m-cresol, o-cresol, p-cresol and 3,4-dimethylphenol; Group 3; methane, pentane, heptane, dichloroethane, 1,2-dichloropropane, propene, toluene, benzene and naphthalene (Park *et al*., 2018) [53]; Group 4; ethene, propene, C4–C10 alkenes, isoprene, allyl alcohol, styrene, cyclopentene, cyclohexene, dihydropyran and indene (Cheung *et al*., 2013; McCarl *et al*., 2018) [54,55]; Group 5; C2–C6 alkanes (Kotani *et al*., 2006) [56]; Group 6; propane, propene, toluene, benzene and naphthalene (Park *et al*., 2018); Group 7; C2–C4 alkenes, 1-hexene, 1-octene, isoprene, acrylic acid, vinyl chloride (VC) and 1,2-cis-dichloroethene (1,2-cis-DCE) (Yang *et al*., 2024) [57]. **(B)** Transcription of each SDiMO group in MAGs and unbinned contigs, colored by SDiMO grouping and split by phylum. Circle area gives summed transcript abundance (TPM); fill gives SDiMO grouping, matching the colors of panel A. Data derived from single field library (WIL2 Before; WBe15_8).

### Mobile genetic element carries a silent, mobilizable putative CO dehydrogenase operon

Most MGEs were transcriptionally active and their transcription tracked that of their predicted hosts (Supplementary Results S3, **Supp. Figure 6**). Screening the MGEs revealed that one 104.4-kb plasmid (WILB_D3 contig_507), assigned to *Nocardioides* sp024703895, carried a single contiguous operon of all three CO dehydrogenase subunits (*coxMSL*) (**Supp. Figure 7**). The CoxL subunit showed highest similarity to *Aeropyrum pernix* (37% amino acid identity over 811 aa) within the marker gene database, falling well below our 60% threshold and thus represents a divergent variant (**Supp. Figure 8**). Aside from low-level transcription of *coxS*, the operon was transcriptionally silent under field conditions (**Supp. Figure 7**), reflecting mobilizable genetic potential rather than active CO oxidation. Although *Nocardioides* species are known to harbor linear megaplasmids encoding metabolic genes [58,59], plasmid-borne CO dehydrogenase has, to our knowledge, not previously been reported in this genus. Similar mobilizable chemolithoautotrophy modules occur in other carboxydotrophs, in *Afipia* (formerly *Oligotropha*) *carboxidovorans*, the *cox, cbb* and *hox* clusters are co-located on the 133-kb megaplasmid pHCG3 adjacent to a conjugal transfer region [60]. Together with recent evidence that CO dehydrogenases in cold soils are frequently plasmid-associated and horizontally transferred [61], this raises the possibility that horizontal gene transfer allows pioneer communities to maintain metabolic potential without individual populations incurring continuous expression costs.

### Acetylene perturbation inhibits methanotrophy with no detectable compensatory trace-gas oxidation

To test for metabolic flexibility under enzymatic inhibition and evaluate surface-subsurface coupling, we applied acetylene (C_2_H_2_) perturbations *in situ* across two field campaigns (2024 and 2025) and in laboratory mesocosms (**Figure 6A, B**). Acetylene is a well-characterized inhibitor that irreversibly blocks the active site of pMMO [62,63], but it is not selective for methanotrophs. As a promiscuous inhibitor of metalloenzymes, it may also target [NiFe]-hydrogenases [64,65], nitrogenases [66,67], and, at least partially, CO dehydrogenases [68]. We therefore treat these trials as a broad-spectrum perturbation of the trace-gas functional network.

**Figure 6.**
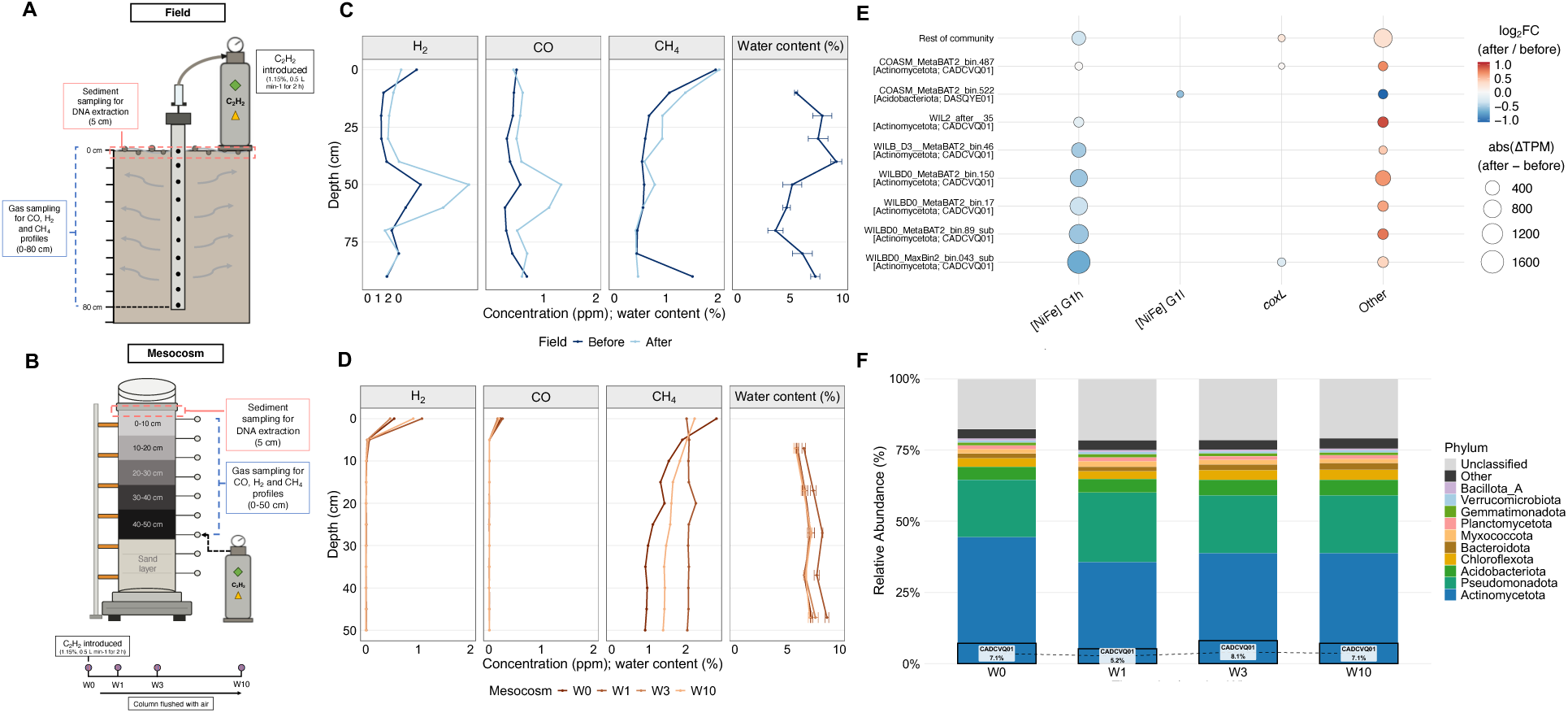
Acetylene perturbation inhibits methanotrophy with no detectable compensatory trace-gas oxidation. **(A)** Design of the field inhibition experiments (1.15% C2H2 in air, 0.5 L min^-1^, 2 h). Field treatments were applied in 2024 at sites C and F and repeated in 2025 at site B with paired sampling for nucleic acids immediately before and 24 h after treatment. **(B)** Design of mesocosm experiments; gas and DNA profiles were collected at week 0, 1, 3, and 10. **(C)** Depth profiles of H2, CO and CH4 before and after treatment with acetylene, and water content (%), in the 2025 field campaign and **(D)** across the mesocosm time series. **(E)** Change in transcription (ΔTPM) after acetylene treatment in CADCVQ01 MAGs, specifically Group 1h and Group 1l hydrogenases and *coxL*, and the contribution to TPM change by the remaining community. Other denotes all other marker genes surveyed. **(F)** Phylum-level community composition in mesocosm surface sediment over ten weeks (metagenomic read-based, Kraken2/Bracken); highlighted is CADCVQ01 which accounted for 7.10% at both week 0 and week 10.

At both process and community level, the forefield microbiome showed no evidence of substrate reallocation following inhibition. In 2024 field trials at sites C and F, acetylene inhibited methanotrophy and suppressed H_2_ oxidation (**Supp. Figure 9,** Supplementary Results S4). Methanotrophy was similarly inhibited in mesocosm columns packed with soil from site B, although H_2_ and CO consumption persisted to low-ppb minima in the top 5 cm regardless of headspace supply (**Figure 6D**). Community structure remained stable across the 10-week incubation, with the dominant hydrogenotrophic lineage CADCVQ01 remaining at 7.10% (**Figure 6F**). The loss of methanotrophy therefore triggered no compensatory shift toward H_2_ or CO oxidation in the mesocosm. In the 2025 field treatment at site B, CH_4_ uptake 24 h after acetylene addition was reduced but not abolished, while H_2_ and CO accumulated at 40-60 cm (**Figure 6C**), i.e., inhibition was only partial. Exploratory metatranscriptomes collected before and 24 h after this 2025 field treatment show the same absence of switching at the transcriptional level (**Figure 6E, Supp. Figure 10**). The largest signal was a severe decline in G1h hydrogenase transcripts, concentrated in the same CADCVQ01 populations that dominate baseline hydrogenase expression: one CADCVQ01 MAG accounted for 32.7% of the total trace-gas transcript decline, and three further CADCVQ01 MAGs for most of the remainder (**Figure 6E**). Notably, *coxL* was not upregulated in response to hydrogenase suppression, it instead declined moderately, again mainly in CADCVQ01, and in those genomes encoding both enzymes *coxL* remained 20- to 35-fold below G1h hydrogenase levels before treatment. This muted response may in part reflect partial inhibition of CO dehydrogenase by acetylene [68].

## Discussion

Forefield soils of the Wildstrubel glacier acted as consistent sinks for atmospheric H_2_ and CH_4_, and although fluxes measured in the field suggested it to be a net source for CO, kinetic assays *ex situ* revealed high-affinity scavenging for all three gases. Whole-soil apparent K_m_ values were in the nanomolar range for both H_2_ (191 nM) and CO (52 nM), comparable to the purified Huc (129 nM) and Mo-CO dehydrogenase (139 nM), despite soil diffusion typically inflating apparent K_m_ [69,70]. Previous work indicates that these energy pathways overlap along the chronosequence rather than replacing one another: sulfur oxidizers dominate in the first years after deglaciation [13], atmospheric H_2_ oxidation proceeds across whole chronosequences [12] and methanotrophy and nitrification strengthen with soil age [15,71].

Despite broad sink capacity and widespread genomic potential, with 57% of recovered MAGs encoding trace-gas machinery, transcriptional activity was narrow. Expression was dominated by G1h [NiFe]-hydrogenases within a few Actinomycetota lineages, namely CADCVQ01 belonging to Thermoleophilia, a class abundant in cold oligotrophic soils that has, to our knowledge, not been validated as an atmospheric H_2_ oxidizer in culture. This pattern provides *in situ* confirmation that G1h [NiFe]-hydrogenases are the primary mediators of atmospheric H_2_ scavenging during carbon-limited persistence [72,73], likely because H_2_ offers a continuously available, highly diffusible atmospheric energy source [74]. In contrast, *pmoA* was absent from MAGs and minimally transcribed in unbinned contigs, and *coxL* was transcribed 20-to 35-fold less than [NiFe]-hydrogenases within MAGs. Therefore, transcript ratios did not mirror process rates, our kinetic assays indicated that CO is oxidized at rates of the same order as H_2_ at ambient concentrations, however *coxL* transcripts were far less abundant than hydrogenase transcripts, both within CADCVQ01 and across the community. This suggests that transcript abundance underrepresents CO oxidation in these soils.

*SmoX* (encoding SDiMOs) ranked second among transcribed markers across generalists including *Pseudonocardia*, JACCQS01, and CADCVQ01. Its homology to C_2_-C_6_ alkane monooxygenases induced by ethane and propane in *Mycobacterium* and *Pseudonocardia* [56] alongside active SDiMO-mediated alkane oxidation at terrestrial gas seeps [75] indicates that short-chain alkanotrophy may represent an unappreciated subsurface energy source complementing atmospheric H_2_ scavenging. Because Wildstrubel sediments entrap thermogenic ethane and propane [21] that is released on disturbance [76], we hypothesize that calcareous forefields on hydrocarbon-bearing bedrock could act as low-flux gas seeps, which remains to be tested with alkane concentration and flux measurements. MGEs may provide an additional layer of metabolic versatility in these pioneer soils. Unlike most MGEs recovered, a 104-kb *Nocardioides* plasmid, carrying a complete but divergent *coxMSL* operon (37% CoxL amino acid identity to the closest reference), was transcriptionally silent, paralleling plasmid-borne chemolithoautotrophy modules in other carboxydotrophs [60,61]. This may represent a mobilizable reserve expressed only under conditions our sampling did not capture, although whether this CoxL is a functional CO dehydrogenase remains unknown.

Perturbation of these communities with acetylene did not reveal detectable metabolic reallocation either on a process, community or expression level in field (sites C and F in 2024; site B in 2025) or mesocosm experiments. At site B in 2025, G1h [NiFe]-hydrogenase transcript abundance in CADCVQ01 was lower 24 hours after treatment without an increase in *coxL,* and eliminating methanotrophy over 10 weeks in mesocosms neither stimulated greater H_2_ or CO uptake nor altered community composition. However, three experimental caveats apply: first, acetylene can inhibit [NiFe]-hydrogenases and partially inhibit CO dehydrogenases [64,65,68], potentially blocking compensation; second, in the active topsoil populations sampled we did not detect methanotrophs and thus we saw no inhibited pathway to switch from; and third, mesocosm topsoil H_2_ and CO were already drawn down to supply-limited, low-ppb levels. The absence of induced switching therefore does not contradict the metabolic flexibility observed in cultured aerotrophs and the presence of multiple aerotrophic genes within single organisms previously seen at glacier forefields [8,11,12,17,18]. In fact, CADCVQ01 populations co-transcribed [NiFe]-hydrogenases, *coxL*, and in some cases RuBisCO, aligning with evidence that high-affinity [NiFe]-hydrogenases and CO dehydrogenases are regulated by internal energetic demands rather than external substrate availability [6,77].

The strong transcriptional signal for hydrogenotrophy is consistent with vertical partitioning across depth, which could reconcile minimal topsoil *pmoA* transcription with strong net CH_4_ uptake. In diffusion-limited soils, high-affinity H_2_ and CO uptake is confined to the uppermost few centimeters [78,79], whereas CH_4_ oxidation in forefield soils peaks at 10-40 cm, where moisture is sufficient to prevent desiccation without restricting diffusion [14,71,80,81]. Our 5 cm samples therefore captured the hydrogenotrophic but not the methanotrophic layer. Across the chronosequence, soil moisture is a likely primary physical driver of gas transport: elevated moisture in wetter, poorly drained younger terrain may restrict diffusion and limit net H_2_ and CH_4_ uptake, even though *ex situ* H_2_ oxidation potential is invariant across the Griessfirn chronosequence [12]. In contrast, well-drained older soils facilitate deeper gas diffusion, maximizing H_2_ and CH_4_ scavenging, while biological CH_4_ capacity grows via a two-order-of-magnitude rise in *pmoA* copy numbers in other Alpine forefields [71], typical of slow-growing high-affinity methanotrophs. CO is decoupled differently: older soils were net CO sources, yet their topsoil sediments scavenged CO with nanomolar affinity, and repacked mesocosms, lacking a subsurface source, drew CO down to low-ppb levels, indicating that a subsurface CO source of yet unknown origin, is large enough *in situ* to reverse the net flux sign. The contrasting efficacy of acetylene across campaigns may reflect how sediment structure and moisture govern gas transport: whereas denser, wetter terrace soils in 2024 (sites C and F) and repacked columns retained inhibitor long enough to abolish methanotrophy, coarse, well-drained deposits at site B in 2025 promoted rapid gas dispersion and only partial inhibition, accompanied by H_2_ and CO accumulation at 40-60 cm. This accumulation of H_2_ and CO at site B might indicate additional, unquantified subsurface sources, e.g., fermentative H_2_ production in moist anoxic microsites, and abiotic CO production or release of gas entrapped in sediment and rock [21,48,82]. By regulating atmospheric gas diffusion, sustaining micro- anoxic niches, controlling for cell physiological stress, and modulating rock-surface degassing through water-table dynamics [19–21,83], soil moisture likely structures this vertical coupling.

Several methodological limitations warrant consideration. Our transcriptomic interpretations rely on two unreplicated libraries from a single site and depth, one collected before and one 24 h after partial acetylene inhibition, alongside homology-based assignments for *smoX* and the plasmid-borne *coxL*. Furthermore, because acetylene inhibits enzyme activity rather than transcription directly, and we lacked an acetylene-free control to distinguish inhibition from physical flushing by the ∼60 L gas injection, field responses cannot be attributed unambiguously to biological processes. Future work should pair replicated, depth-resolved metatranscriptomics with isotope-tracer estimates of gross H_2_, CH_4_ and CO turnover to resolve active layers and masked uptake, alongside alkane profiling, ^13^C-alkane stable-isotope probing, and biochemical characterization of forefield SDiMOs.

In conclusion, pioneer forefield communities encode broad potential for trace-gas oxidation yet transcribe a narrow repertoire *in situ*, concentrating surface metabolism on atmospheric H_2_ with transcriptional evidence for the use of short-chain alkanes of possible subsurface origin. We detected no induced pathways switching after perturbation, although our design cannot exclude it. These communities’ function within a vertically structured column governed by atmospheric inputs from above, geological gas subsidies from below, and soil moisture dynamics in between. Ultimately, surface omics and net fluxes cannot be interpreted in isolation: vertical partitioning can decouple topsoil scavenging from deeper methanotrophy and cause net surface fluxes, particularly for CO, to underestimate gross uptake.

## Data availability

Oxford Nanopore metagenomic reads, Illumina metatranscriptomic reads and high-quality MAGs have been deposited in the NCBI SRA and Genomes under BioProject PRJNA1559487. Remaining medium-quality MAGs, the *Nocardioides* host-associated plasmid containing *coxMSL* and phylogenetic trees of SDiMOs, *amoA/pmoA* and *coxL* are available at Zenodo under https://zenodo.org/records/24159710.

## Code availability

Analysis scripts are available at https://github.com/envmicrobiol-ethz/Herbert_et_al_2026; software versions are in the Methods.

## Author contributions

Josephine Herbert (Formal analysis, Investigation, Writing—original draft, Writing—review & editing), Bledina Dede (Formal analysis, Investigation, Writing—original draft, Writing—review & editing), Lorenz Koschitz (Investigation, Resources, Formal analysis), Julian Sarbach (Investigation, Resources, Formal analysis), Martin H. Schroth (Conceptualization, Formal analysis, Investigation, Project administration, Resources, Writing—review & editing), and Marie C. Schoelmerich (Conceptualization, Formal analysis, Funding acquisition, Investigation, Project administration, Resources, Writing—original draft, Writing—review & editing).

## Acknowledgements

We thank Lukas Wanner for technical support and Emilie J. Skoog and Luzia Stalder for their helpful discussion regarding metagenomic analyses.

