## Supplementary material for "Depth-partitioned oxidation of atmospheric hydrogen, carbon monoxide and methane in a glacier forefield soil": Supp.

### Supplementary Information

This Supplementary Information file contains Supplementary Figures, Supplementary Methods describing more detail of what is written in the main text, and Supplementary Results supporting the results of the main text. Supplementary Tables are provided as a separate Excel spreadsheet.

### Supplementary Figures

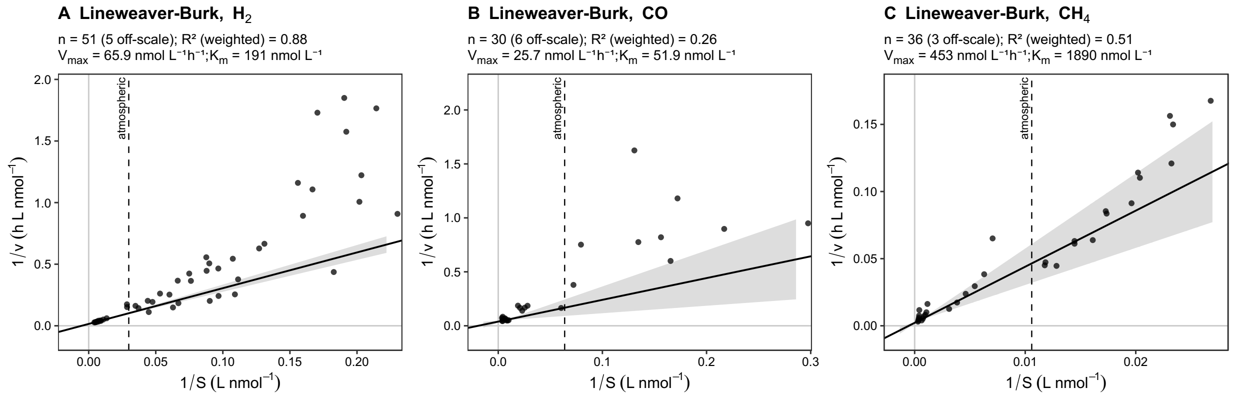


**Supp. Figure 1. Lineweaver-Burk regressions underlying the apparent kinetic parameters.** Double-reciprocal plots for $H_2$ **(A)**, CO **(B)** and $CH_4$ **(C)**, from the incubations plotted in **Figure 2D-F**. Lines give weighted linear regressions using $V^4$ weights; only observations with $1/v < 200$ h g nmol $w^{-1}$ are plotted and fitted. Each panel gives the number of observations (n), the weighted $R^2$ of the regression, and the resulting $V_{max}$ and $K_m$ (**Supp. Table 16**). These estimates are reported alongside non-linear least-squares fits in **Supp. Table 15** as a cross-check rather than as primary values.

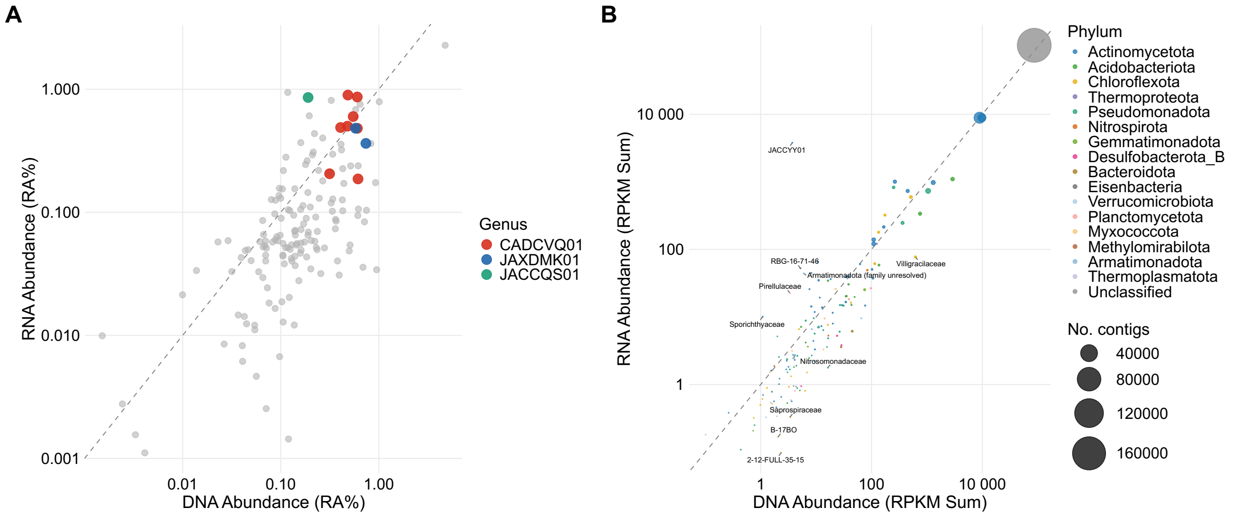


**Supp. Figure 2. MAG and unbinned DNA abundance versus transcriptional activity. (A)** Relative abundance (RA%) of each of the 159 MAGs in the metagenome (DNA) and metatranscriptome (RNA) from the sample before acetylene treatment (WIL2 Before; WBe15_8); the dashed line is 1:1. MAGs of the genera CADCVQ01, JAXDMK01 and JACCQS01 are highlighted. **(B)** Summed RPKM in DNA and RNA for each family of unbinned contigs; point size indicates the number of contigs, color the phylum, and the dashed line is 1:1. Selected families far from the line are labelled.

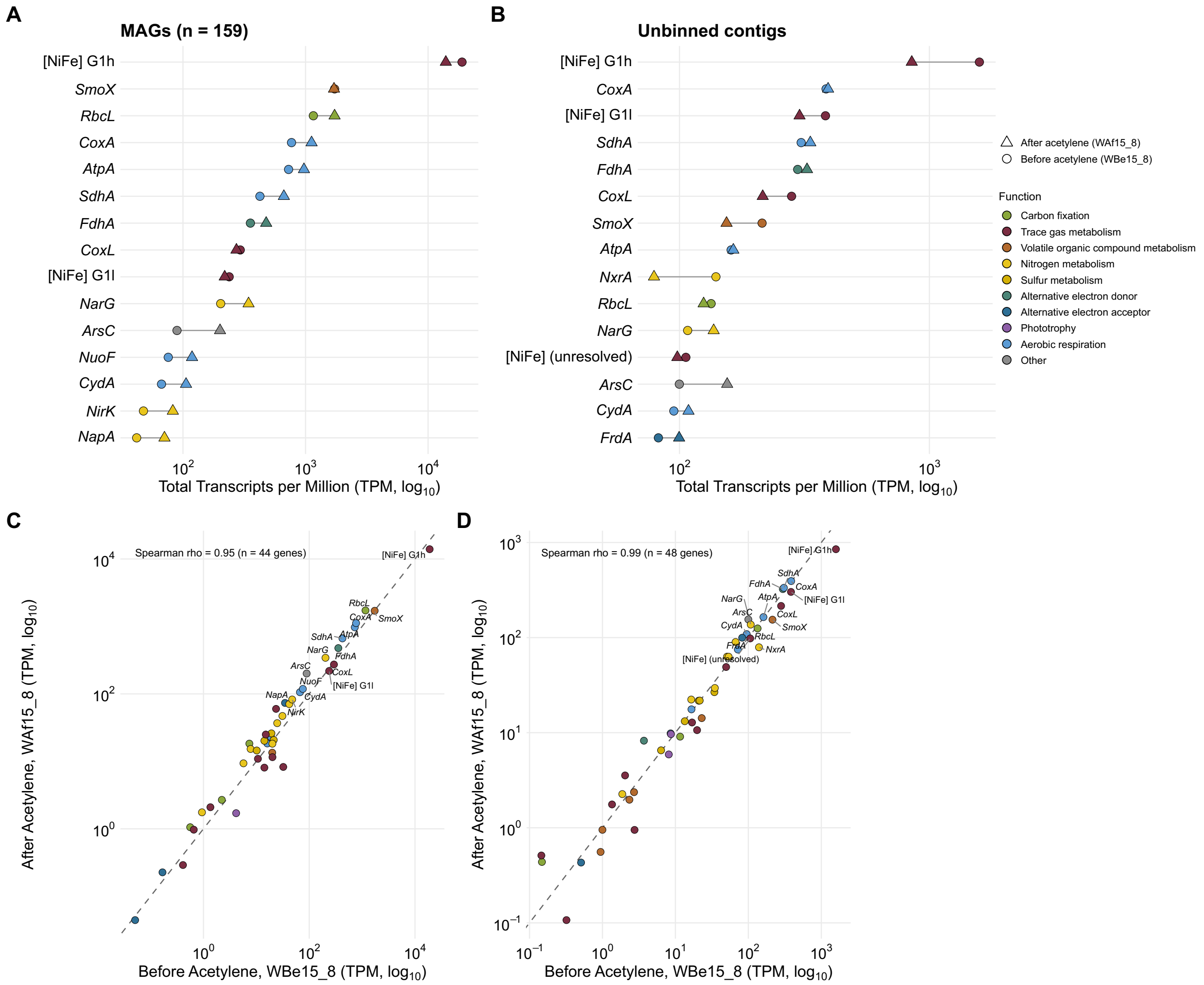


**Supp. Figure 3. Transcription of the top 15 marker genes before and after acetylene treatment. (A, B)** The 15 most highly transcribed marker genes in MAGs **(A)** and unbinned contigs **(B)**, ranked by total transcripts per million (TPM, $\log_{10}$) summed across all genes of that type. Circles show the sample before acetylene treatment (WIL2 Before; WBe15_8) and triangles the sample after (WIL2 After; WAf15_8); color indicates functional category. **(C, D)** Total TPM of every detected marker gene before versus after acetylene treatment for MAGs **(C)** and unbinned contigs **(D)**; the dashed line is 1:1 and the Spearman correlation is shown (MAGs; $\rho = 0.95$, n = 44 genes; unbinned; $\rho = 0.99$, n = 48 genes). Labelled points are the top 15 genes from **(A, B)** with the same coloring.

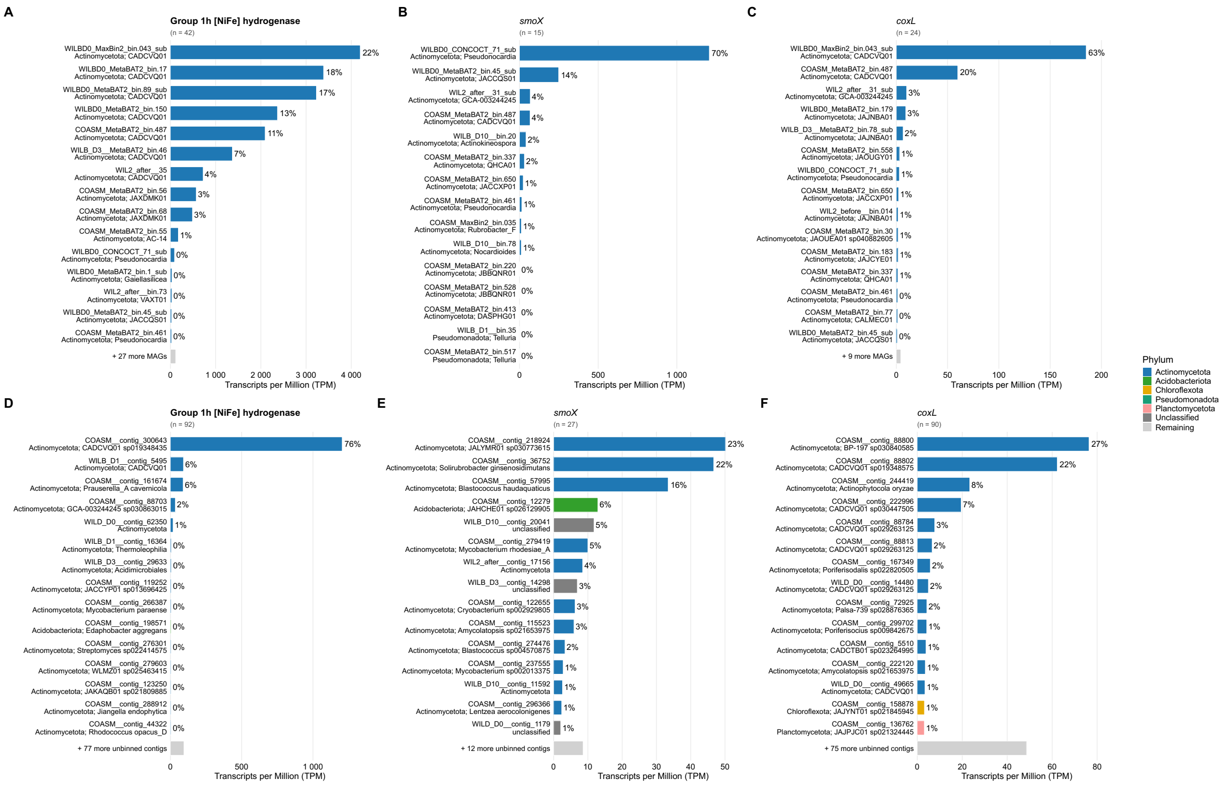


**Supp. Figure 4. Contributors to Group 1h hydrogenase, *smoX* and *coxL* transcription.** The 15 highest-transcribing MAGs **(A-C)** and unbinned contigs **(D-F)** for Group 1h [NiFe] hydrogenase **(A, D)**, *smoX* **(B, E)** and *coxL* **(C, F)**, ranked by transcripts per million (TPM) in the sample before acetylene. Percentages show each MAG's or contig's share of the total TPM for that gene; n is the number of MAGs or contigs carrying the gene, and the grey bar pools the rest. Bars are colored by phylum, and labels give the lowest assigned taxon.

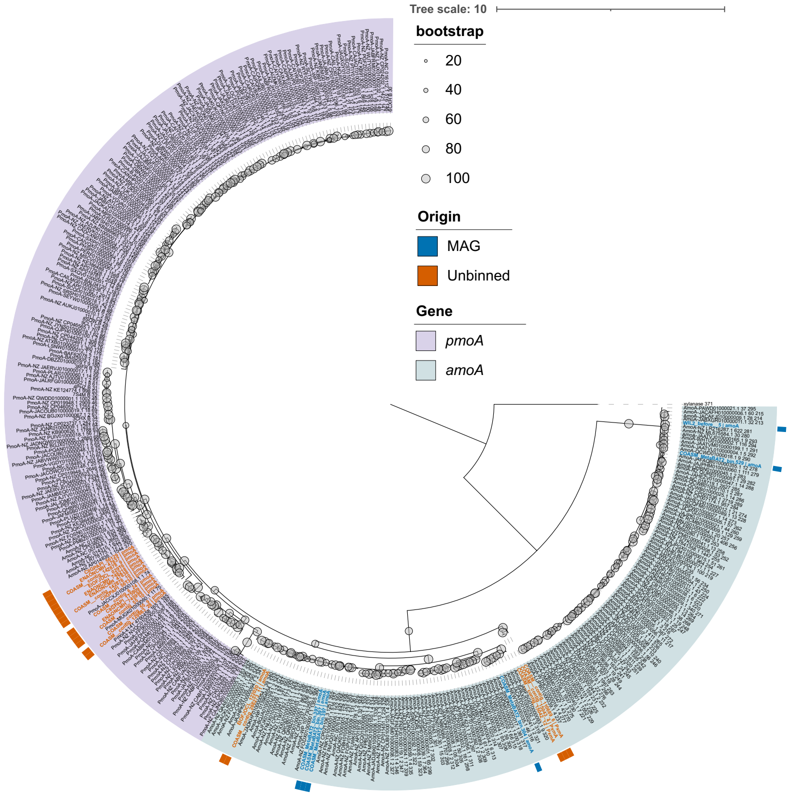


**Supp. Figure 5. Discrimination of particulate methane monooxygenase from ammonia monooxygenase.** Phylogenetic tree of all genes recovered in as particulate methane monooxygenase protein A (*pmoA*) or ammonia monooxygenase (*amoA*) from MAGs (n=159) and unbinned contigs. Highlighted in blue are proteins from MAGs and in orange from unbinned contigs.

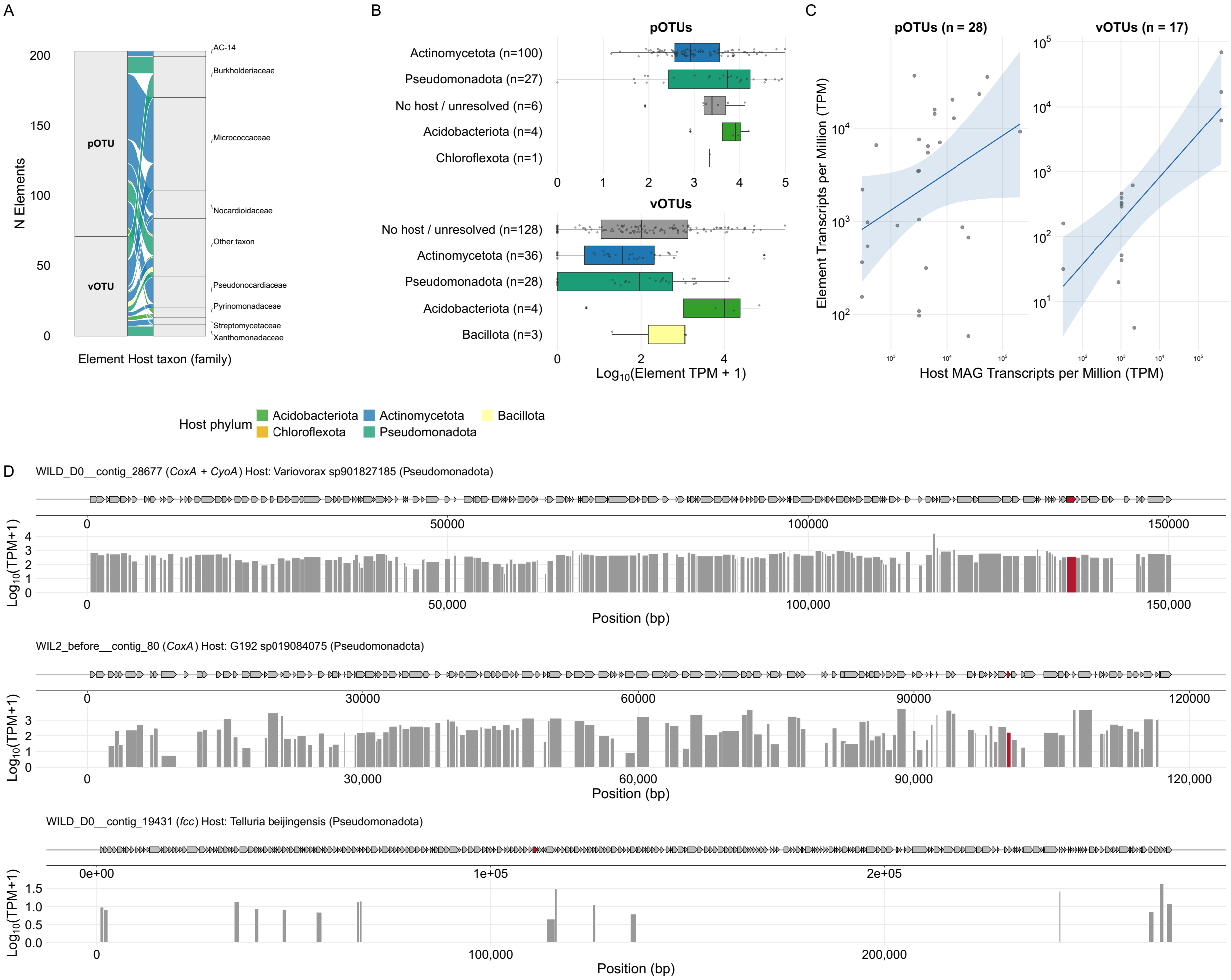


**Supp. Figure 6. Extrachromosomal elements are transcriptionally active and carry mobilizable metabolic capacity. (A).** Predicted hosts of plasmid-associated (pOTU, n = 138) and virus-associated (vOTU, n = 199) elements at family level (iPHoP, BLAST method, >= 90% confidence tier), colored by host phylum. Elements without a confident prediction are shown as “unknown host”. **(B)** Cargo-carrying elements, drawn as +/- kb windows around each locus at a common scale; hatched ends denote omitted sequence. Important metabolic genes are colored, other coding sequences in grey. The track beneath each element gives transcript coverage ($\log_{10}$(TPM+1)) in the sample before acetylene treatment (WIL2 Before; WBe15_8). **(C)** Transcript abundance of each element against that of its predicted host MAG, for pOTUs (Spearman’s ρ = 0.38 with n = 28, p = 0.044) and vOTUs (ρ = 0.72 with n = 17, p = 0.001) in WBe15_8. **(D)** For each pOTU carrying a marker gene, the upper track shows the predicted genes (arrows show direction of transcription) and the lower track shows gene transcription as $\log_{10}$(TPM+1) before acetylene. Marker genes are in red: *coxA* and *cyoA* (WILD_D0__contig_28677), *coxA* (WIL2_before__contig_80) and *fcc* (WILD_D0__contig_19431).

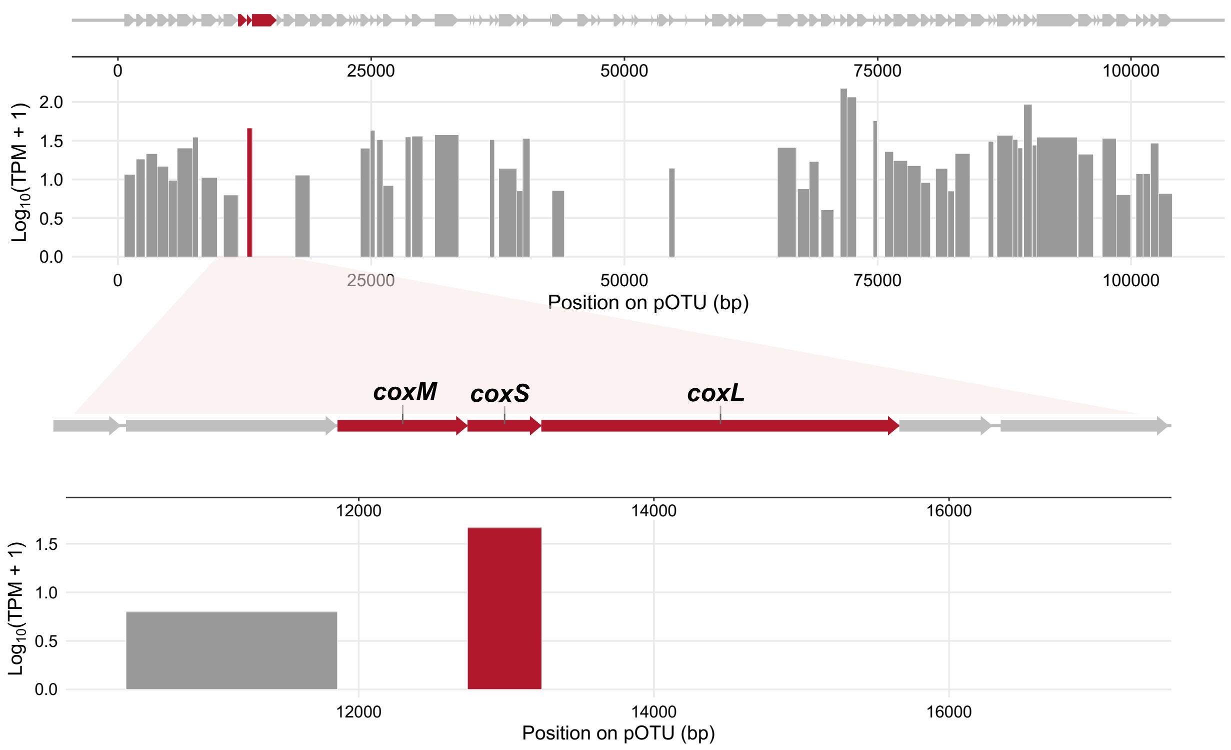


**Supp. Figure 7. Transcription of *coxMSL* operon found in the pOTU host-linked to *Nocardioides*.** Top: gene map and per-gene transcription ($\log_{10}$(TPM+1)) across the full ~104-kb plasmid WILB_D3__contig_507 (*Nocardioides* sp024703895), with the *coxMSL* operon highlighted in red. Bottom: zoom on the operon and its neighboring genes, once again showing per-gene transcription ($\log_{10}$(TPM+1)) in the sample before acetylene treatment (WIL2 Before; WBe15_8).

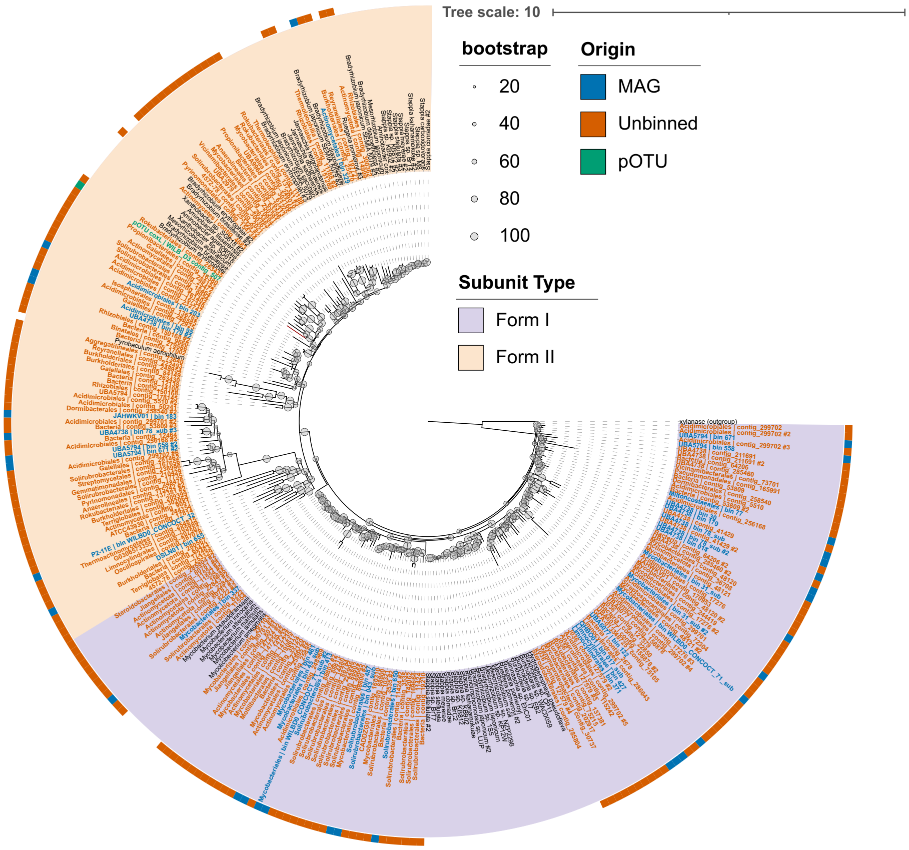


**Supp. Figure 8. Phylogeny of *coxL* recovered from MAGs, unbinned contigs and pOTU.** Phylogenetic tree of all genes recovered as *coxL* from MAGs (n=159) and unbinned contigs, including the *coxL* identified as a part of an operon within the 104.4 kb plasmid (WILB_D3__contig_507). Highlighted in blue are proteins from MAGs and in orange from unbinned contigs and in green the *coxL* identified from the pOTU.

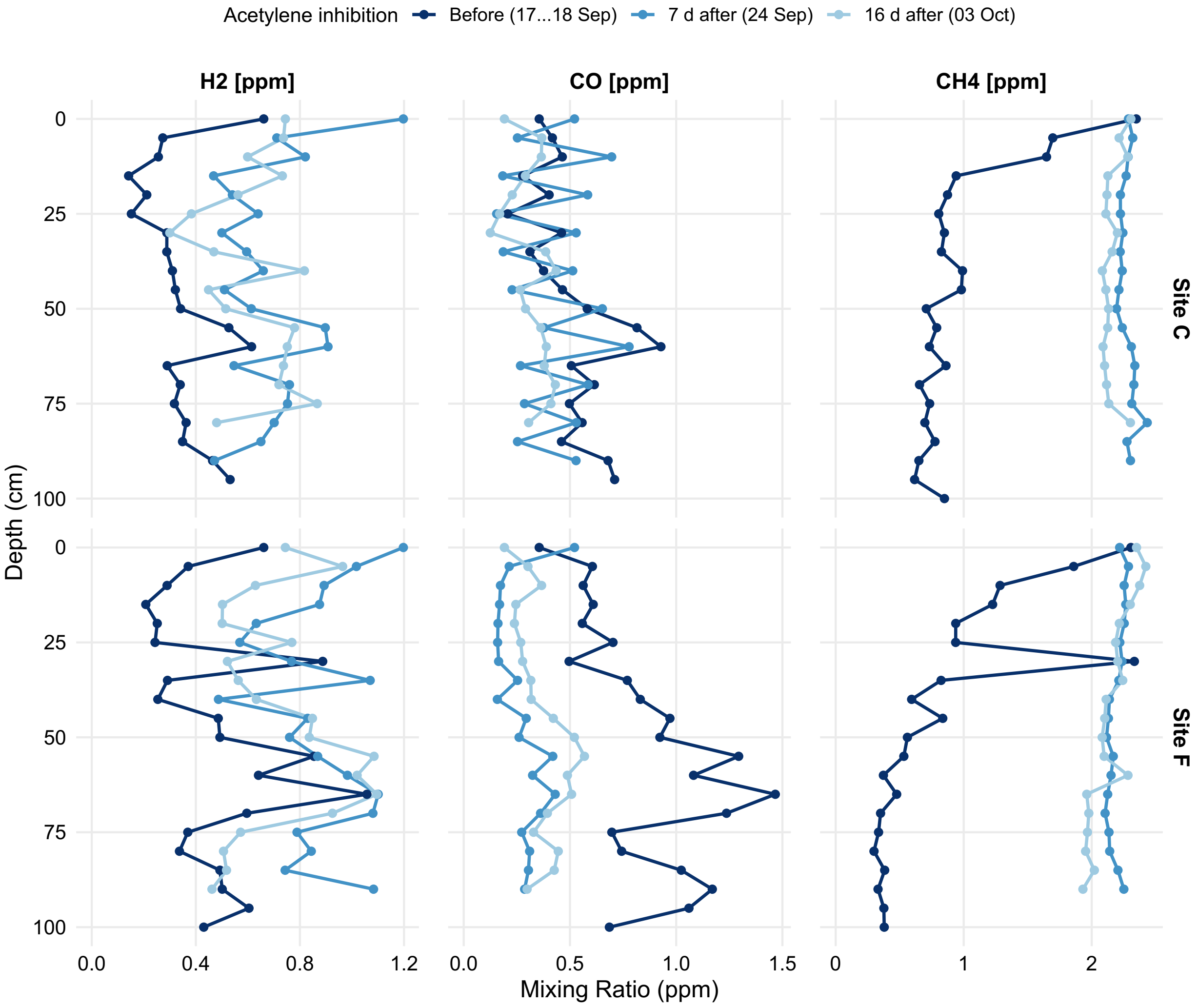


**Supp. Figure 9. Field and mesocosm column inhibition experiments vertical gas profiles from 2024.** Depth profiles of $H_2$, CO and $CH_4$ mixing ratios (ppm) at two field sites (C and F) before (17-18 Sep), 7 d after (24 Sep) and 16 d after (3 Oct) acetylene addition conducted at sites C and F (**Supp. Table 19**).

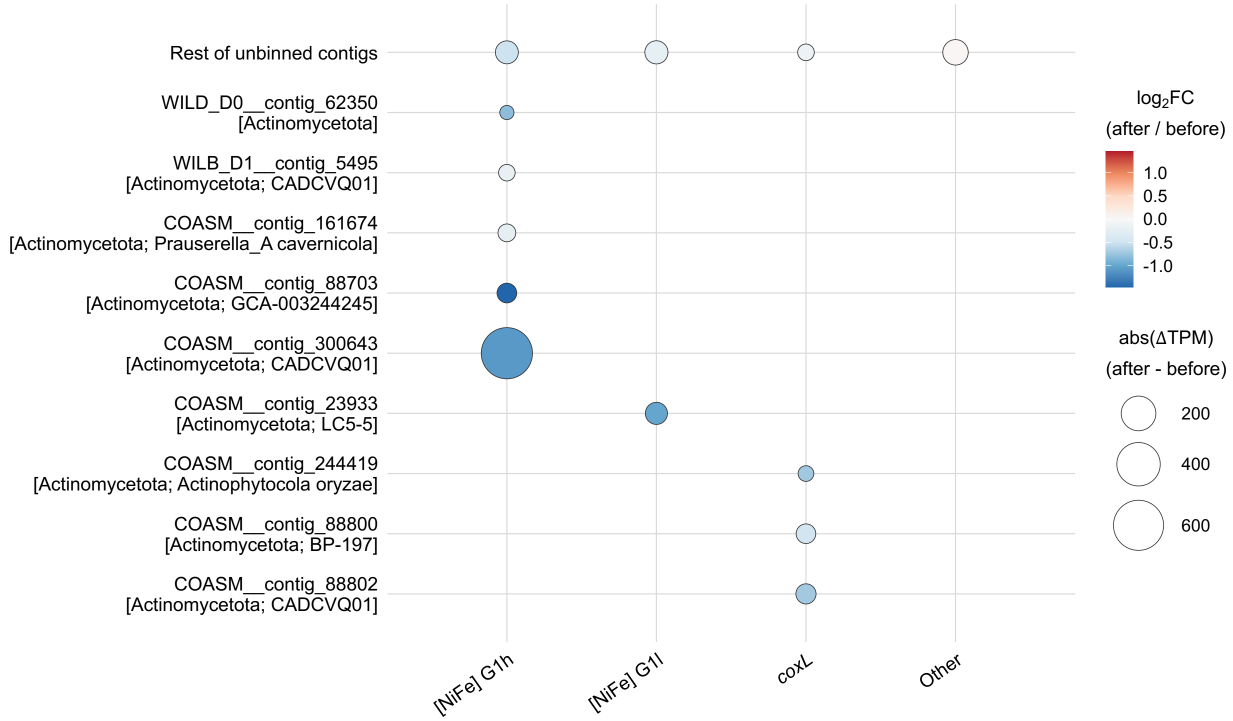


**Supp. Figure 10. Response to acetylene inhibition across unbinned contigs.** Change in transcription (ΔTPM) after acetylene treatment in unbinned contigs, showing top 9 largest change in specifically Group 1h and Group 1l [NiFe]-hydrogenases and *coxL*, and the contribution to TPM change by the remaining community. Other denotes all other marker genes surveyed. Symbol area gives absolute change in transcript abundance (ΔTPM); fill gives $\log_2$ fold change (FC), with warm colors denoting higher abundance after treatment.

### Supplementary Methods

#### S1. Field site and sampling campaigns

In August to October 2024, soil-gas and water-content profiles were measured at ten locations spanning a young (~20 years post-deglaciation) and an older (~50 years post-deglaciation) forefield section, and acetylene inhibition experiments were performed at sites C and F, without taking samples for molecular analyses.

#### S2. Soil gas and water-content profiles

Soil gas profiles were measured at 10 cm depth intervals using the poly-use multi-level sampling system, as described by Nauer *et al.* (2012), from permanently installed perforated access tubes left to consolidate for two weeks before first sampling. $CH_4$ samples were stored in $N_2$-flushed glass vials with butyl stoppers. $H_2$ and CO samples in Ar-flushed, evacuated 10 mL Exetainer vials modified with silicone plugs, steel bolts and O-rings to limit contamination and diffusive gas exchange [1]. Vial pressures were recorded before and after injection to correct for dilution, and residual vial contamination was quantified with control vials per campaign. Volumetric water content was measured at each depth in triplicate with a capacitance profile probe (PR2, Delta-T Devices); at three sites, iButton loggers (DS1921G) recorded temperature at 10 cm intervals every 2 h.

#### S3. Soil-atmosphere flux estimation

Fluxes for $CH_4$ and $H_2$ in the field and in columns were calculated by fitting a steady-state diffusion-reaction model (**eq. 1,** [2,3]) to the respective gas concentration profiles. For this, the effective diffusion coefficient in **eq. 1** was independently derived from air-filled porosity and soil tortuosity (determined from water content measurements using the WLR-Marshall model [4], with total porosity of $\theta_t$ = 0.42 [1]), as well as temperature-adjusted free-air diffusion coefficients [5] using iButton field temperature data. Net soil-atmosphere fluxes ($J_{net}$) were then computed from the gradients of the modeled concentration profiles evaluated at z = 0 (**eq. 2**) using Fick's first law of diffusion (**eq. 3**). For $CH_4$, entire soil concentration profiles were used in fitting field and column data. For $H_2$, entire concentration profiles were used in fitting column data, whereas fitting of field data was limited for consistency to depths of 0 to 30 cm, as some $H_2$ concentration profiles at depths > 30 cm failed to follow the general form of the diffusion-reaction model.

This issue was also noticeable for CO concentration profiles, which failed to fit the general form of the diffusion-reaction model for both field and column data sets. Thus, instead of **eqs. 1** and **2**, linear regression was employed to obtain CO concentration gradients at $z$ = 0, which were then used to calculate CO fluxes using **eq. 3**. For all flux calculations, a positive $J_{net}$ value indicates net gas flux into the soil (uptake) and vice versa. Parameter and flux uncertainties were estimated from the model fits and propagated with the measurement errors of porosity and gas concentration.

$$C_z = (C_0 - C_\infty)\exp\left(-z/\sqrt{\frac{D_{eff}}{\theta_a k}}\right) \quad \textbf{(eq. 1)}$$

$$\frac{dC_z}{dz}_{z=0} = -\sqrt{\frac{D_{eff}}{\theta_a k}}\,(C_0 - C_\infty) \quad \textbf{(eq. 2)}$$

$$J_{net} = -D_{eff}\frac{dC_z}{dz}_{z=0} \quad \textbf{(eq. 3)}$$

where $C_z$ is the gas concentration at depth $z$ (cm); $C_0$ is the gas concentration at the soil surface ($z = 0$); $C_\infty$ is the fitted threshold gas concentration at depth; $D_{\text{eff}}$ is the effective soil diffusion coefficient ($cm^2\ s^{-1}$); $\theta_a$ is the air-filled porosity ($cm^3\ cm^{-3}$); and $k$ is the first-order consumption rate constant ($s^{-1}$).

#### S4. Mesocosm column construction

Approximately 20 kg of sediment was collected by shovel and hand adjacent to site B for inclusion in mesocosm experiments. To preserve the vertical structure, approximately 2 kg of sediment was collected per 10 cm depth interval to a depth of 50 cm. The sediment was sieved to 8 mm and stored at ambient temperature. After ~7 h on arrival at the laboratory, bags were opened to allow air exchange and stored at 15 °C in the dark for three days until the sediment was filled into the columns.

A ceramic plate was placed at the bottom of the Plexiglas columns (diameter; 11 cm, depth; 80 cm, located in a 15 °C climate chamber) to support the porous media, followed by a 20 cm layer of quartz sand. The sediment was filled depth-by-depth to preserve vertical stratification, and each layer was consolidated by tapping. The columns were equipped with ports of stainless-steel tubes connected to external 3-way valves every 5 cm allowing for gas sampling without sediment disturbance, and time-domain-reflectometry (TDR) water content sensors (10 cm long, three-prong TDR probes connected to a TDR100, Campbell Scientific, Loughborough, UK) every 10 cm. Column headspaces were continuously flushed with compressed air through a lid valve, and headspace $H_2$ and CO concentrations were included as covariates in all statistical models. Three weeks after packing prior to inhibition, 360 mL of rainwater was added to the top of the columns, simulating a ~40 mm precipitation event.

#### S5. Acetylene treatment experiments

Acetylene (1.15 vol% in air) contained in a pressurized cylinder (60 L, 34 bar; ALL IN GAS, Starnberg, Germany) served as inhibitor source in all experiments.

**Field, 2024 (sites C and F).** The ground surface around the access tube (AT) was covered with plastic foil and weighted with rocks and sediment; at site C, a steel cylinder (~30 cm diameter) was additionally inserted 47 cm into the ground to confine lateral gas spread. The AT was closed with a rubber plug fitted with a Luer connector, and the acetylene mixture was delivered at ~0.5 L $min^{-1}$ for 2 h. Soil-gas profiles were measured immediately before and after treatment (7 and 16 days).

**Field, 2025 (site B).** The same delivery setup was used, although with no steel cylinder, with soil-gas profiles measured and sediment sampled for nucleic acids (5 and 15 cm; although the deeper depth failed nucleic acid extraction) immediately before and 24 h after treatment.

**Columns.** The acetylene mixture was delivered through the lowest sampling port into the basal quartz sand at 0.5 L $min^{-1}$ for 2 h with three valves opened in the lid, followed by flushing with synthetic air for 15 min. Sediment for DNA-based community profiling was collected from the top 5 cm at weeks 0, 1, 3, and 10 and stored at −80 °C; gas and water-content profiles were measured at the same time points.

#### S6. Apparent uptake kinetics

Apparent Michaelis-Menten parameters for $H_2$ and CO consumption were estimated in 117 mL serum bottles containing ~30 g of sediment adjusted to ~10% water content, incubated at 15 °C. Headspace concentrations were elevated to ~3 or ~6 ppm and their decline followed over 5-7 h on three days, yielding consumption rates across ~60-7,000 ppb. CO treatments were sealed with crimp-cap septa pierced once by a fixed needle/three-way-valve assembly. Rates were computed as local slopes of the concentration-time series and normalized to sediment dry mass. Parameters were estimated by nonlinear Michaelis-Menten curve fitting (nlsMicrobio; [6]) and by weighted Lineweaver-Burk regression (weights $v^4$), the latter yielding narrower confidence intervals and reported in the main text; specific affinities were calculated as $a^o_s = V_{max(app)}/K_{m(app)}$ [7,8].

Apparent $CH_4$ uptake kinetics were studied in the mesocosm column packed with sediment from site B under conditions of ambient and elevated $CH_4$ concentrations in the column headspace. For ambient $CH_4$ concentration, the column headspace was continuously flushed with compressed air (see above). For elevated $CH_4$ concentrations, a 40 L gas bag was filled with hydrocarbon-free air (hydrocarbon-free synthetic air, Linde Gas AG, Zurich) amended with $CH_4$ from standards of known concentration (10 and 100 ppm $CH_4$ in synthetic air, Messer Schweiz AG, Lenzburg) to yield final $CH_4$ concentrations of ~ 2, 5, 10, 50, and 100 µL/L. The gas bag was then connected to the column headspace, and a gas-tight pump was used to continuously flush the column headspace with the respective gas mixture at a flow rate of ~ 1.3 L $h^{-1}$. Experiments were performed in sequence from lowest to highest $CH_4$ concentration. After connecting a new gas bag, the column was left to equilibrate for 18 h before collecting gas samples. Apparent kinetic parameters were then calculated by quantifying mean $CH_4$ concentration, flux and turnover rate for each depth interval for each of the $CH_4$ headspace concentrations employed.

#### S7. Gas composition analysis

The Peak Performer 1 RCP (910 - Series) by PEAK Laboratories (PEAK LABORATORIES, LLC, California, USA) was used to analyze the $H_2$ and CO gas samples. The gas analyzer was initially calibrated for $H_2$ using a linear two-point calibration in the range of 250 to 5000 ppb and for CO using a linear one-point calibration in the range of 0 to 500 ppb. These calibrations were done using reference gas mixtures of known concentrations. Subsequently, the concentrations of both gases in a reference air bottle were determined, which was from then on used to calibrate the gas analyzer before each measuring day using a one-point calibration. Usually, the machine took around 10 injections until the measurements were stable within a range of ± 5 ppb. Sample injections were done using a syringe, injecting a volume of 3 mL. $CH_4$ in soil-gas samples was quantified using a gas chromatograph with flame-ionization detector (GC-FID; 8610C, SRI Instruments Europe, Bad Honnef, Germany) equipped with two columns in sequence: a Porapak Q column followed by a Hayesep D column and operated at 60°C, using $N_2$ as carrier gas. The FID was operated at 180°C. A one-point calibration of the GC-FID was done using compressed air (Linde Gas AG, Zurich) with a previously established $CH_4$ concentration of 1.98 ± 0.04 ppm, by measuring 3 replicas and fitting a calibration curve through the origin.

#### S8. Statistical analyses

Treatment and covariate effects on fluxes were assessed in R with linear and multiple linear regression and two-way ANOVA. Differences in flux between locations and sampling dates were tested with a two-way ANOVA. The effects of ambient (or headspace) gas concentration and soil water content on fluxes were tested by multiple linear regression. The effect of acetylene was tested by regression of flux on treatment state (before or after acetylene), with and without ambient concentration as a covariate, and the models were compared with F-tests. Model assumptions were checked with residual diagnostic plots, Shapiro-Wilk tests for normality of residuals and Levene’s test for homogeneity of variance. Given the unreplicated design of the metatranscriptomic sampling, no inferential statistics were applied to between-library transcript comparisons, which are reported descriptively. Relationships between datasets (for example DNA versus RNA abundance, or transcript abundance before versus after acetylene) are reported as Pearson or Spearman correlation coefficients.

#### S9. DNA and RNA extraction

Total RNA and DNA were co-extracted from 4-5 g of sample using the Qiagen RNeasy PowerSoil Total RNA Kit (Qiagen, Hilden, Germany; Cat. No. 12866-25) with custom modifications. Soil samples were pre-treated with 1.5 mL of a 10 µM dNTP solution, followed by 1.5 mL of lysozyme buffer (10 mM Tris, 1 mM EDTA, 10 mg/mL lysozyme) and incubated at 37 °C for 20 min. Next, 3.5 mL of PowerBead solution, 0.3 mL of Solution SR1, and 0.1 mL of Solution IRS were added, and samples were incubated at 50°C for 1 h with manual shaking for 10 s every 15 min. Phenol/chloroform/isoamyl alcohol (4.5 mL) was added, and samples were vortexed briefly until homogenous, followed by continuous vortexing for 2 min. Subsequent RNA purification and DNA co-elution (using the Qiagen RNeasy DNA Elution Kit (Qiagen, Hilden, Germany; Cat No. 12867-25)) were performed according to the manufacturer's protocols.

#### S10. MAG binning, contig annotation, unbinned contig processing and marker-gene identification

For MAG generation, reads were mapped back to the assemblies using minimap2 v2.22 [9] and samtools v1.16.1 [10]. Contigs were binned using MetaBAT2 v2.17 [11], CONCOCT v1.1.0 [12] and MaxBin2 v2.2.7 [13], and DAS Tool v1.1.7 [14] selected the best bins per assembly. Bin quality was assessed with CheckM2 v1.0.2 [15] applying a minimum completeness of 50% and a maximum contamination of 10% per MIMAG standards [16]. Bins from all assemblies were pooled and dereplicated at 95% ANI using dRep v3.4.5 [17]. Co-assembly bins replaced single-sample bins where they were more complete according to CheckM2 output. Taxonomy was assigned with GTDB-Tk v2.4.0 [18] was employed, using the GTDB r220 database. MAG abundance was estimated with CoverM v0.6.1 using minimap2 in ONT mode and a minimum read identity of 90%.

Unbinned contigs were extracted from all seven per-sample assemblies and the co-assembly by removing contigs assigned to MAGs during binning, identified using the contig-to-MAG mapping tables generated by DAS Tool v1.1.7 [14]. All unbinned contigs were mapped against the dereplicated MAGs using minimap2 v2.22 (--ax asm20) [9] and contigs with ≥80% query coverage against any MAG were excluded. The remaining contigs were dereplicated at 99% nucleotide identity using MMseqs2 v14 (linclust, --min-seq-id 0.99, --cov-mode 1, -c 0.80) [19] to remove redundant sequences and a representative sequence was retained for each cluster.

To taxonomically annotate the assembly-level contigs, DIAMOND v2.1.10 [20] was used to align annotated proteins obtained via Prodigal v2.6.3 [21] to the GTDB r220 database, the lowest common ancestor was then assigned using TaxonKit v0.19.0 [22].

For functional annotation, Prodigal v2.6.3 was used to generate protein profiles. Metabolic marker genes were searched using a curated protein database of 59 metabolic marker genes (https://github.com/GreeningLab/GreeningLab-database; Accessed 09.10.2026). These were searched as described in Ricci *et al*., 2025. Briefly, DIAMOND v2.1.10 was used to search for these genes (including *pmoA*) and then the output filtered using a query coverage of 80% and identity threshold of 50%, except for *rho* (40%), *nuoF, mmoX*, [FeFe]-hydrogenases, *amoA, nxrA, rbcL* (60%), *psaA* (80%), *psbA*, *isoA, atpA, aro, ygfK* (70%), and *hbsT* (75%). Hits were retained only if they were at least 40 amino acids (aa) in length or with at least 80% query or 80% subject coverage. For predicted proteins, the same identity thresholds were used as above except for *atpA* (60%), *psbA* (60%), *rdhA* (45%), *cyc2* (35%), and *rho* (30%) [23–28]. *coxL* was identified using 709 references and cut-offs of alignment length >40 aa AND identity >60% [23].

Hydrogenases were similarly identified using HMMsearch (HMMer v3.4) [29] using the HydDB database (2019) (e-value cut-off: 1e-5) [30], these were then further quality-controlled and classified into groups using DIAMOND v2.1.10 (--max-target-seqs 1) against HydDB and identity cutoffs applied per class (NiFe Group 4 >50%, other NiFe >30%, FeFe >45%, Fe >50%) [30].

#### S11. Phylogenetic tree analysis

Soluble diiron monooxygenase (SDiMO) protein sequences were queried against the clustered non-redundant (nr_clust) database using BLAST [31] to identify homologs. Retrieved sequences were filtered using thresholds of ≥50% sequence identity and ≥70% alignment coverage. Representative SDiMO sequences spanning seven previously described groups [32] were included to contextualize the phylogenetic placement of the recovered sequences. Separately, particulate methane monooxygenase (pmoA subunit) and ammonia monooxygenase (amoA subunit) protein sequences obtained from the Greening database (described above) were used as references to reconstruct a second phylogenetic tree. For carbon monoxide dehydrogenase (coxL subunit), reference sequences representing forms I and II following Dede *et al.*, 2024 [33], were included alongside the recovered genomic, unbinned and plasmid-associated *coxL* sequences to determine their phylogenetic affiliations.

Both sequence datasets were independently aligned using MAFFT v7.505 [34] with the L-INS-i algorithm (--localpair --maxiterate 1000), and poorly aligned regions were removed using trimAl v1.5.rev0 [35] (-automated1). Maximum-likelihood phylogenetic trees were independently inferred using IQ-TREE3 v3.0.1 [36], with the best-fit substitution model selected using ModelFinder (-m MFP). Branch support was assessed using 1,000 ultrafast bootstrap replicates (-bb 1000).

#### S12. MGE identification, annotation and activity analysis

MGEs were identified in the same manner as Dede *et al.*, 2026 [37]. Briefly, geNomad v1.12.0 [38] was used to identify preliminary elements. Viral contigs were quality-filtered using CheckV v1.0.3 [39] (≥3 kb, ≥3 genes) and dereplicated at 95% nucleotide identity with ≥85% coverage using CD-HIT-EST v4.8.1 [40] to define viral OTUs (vOTUs); plasmid sequences were dereplicated using dRep v3.4.5 [17] (95% ANI, 50% coverage) to define plasmid OTUs (pOTUs). Their activity was assessed by using coverM to map the metatranscriptome to the vOTU and pOTU contigs for whole activity (--min-read-percent-identity 90) and then applying the TranscriptM pipeline for resolving marker gene transcription. Host linkage was determined using iPHoP v1.4.1 [41] with MAGs added to the database (June 2025 release, GTDB r220) for closer associations.

### Supplementary Results

#### S1. Proposed naming of the novel genus CADCVQ01

The genus-level lineage CADCVQ01 (family Solirubrobacteraceae, order Solirubrobacterales, class *Thermoleophilia*, phylum Actinomycetota; GTDB r220) has no formally described or named member. We propose the name *Alpihydrogenophilus* gen. nov. (Etymology: L. fem. pl. n. *Alpes*, the Alps; N.L. neut. n. *hydrogenium*, hydrogen; Gr. masc. adj. *philos*, loving; N.L. masc. n., an organism from the Alps that loves hydrogen), with *Alpihydrogenophilus wildstrubelensis* sp. nov. (N.L. masc. adj., pertaining to the Wildstrubel massif, Swiss Alps) as type species. The type of the species is the genome WILBD0_MetaBAT2_bin.150 (isolate code WILMAG04; BioSample [SAMN65124274]; 3.24 Mb in 13 contigs, N50 365 kb, G+C 71.7%, 96.2% complete, 0.63% contaminated by CheckM2 v1.0.2 [15], with one complete 16S-23S-5S rRNA operon), which encodes and transcribes a G1h [NiFe]-hydrogenase and RuBisCO. The genus comprises eight MAGs from the Wildstrubel glacier forefield that represent separate species, only one of which is named here. Genomes of the genus encode a high-affinity group 1h hydrogenase, and the genus is inferred to oxidize atmospheric $H_2$ and is highly active within the topsoil of this forefield, comprising the most active metabolism. The names are registered in the SeqCode Registry (genus: https://seqco.de/i:58162; species: https://seqco.de/i:58165).

#### S2. Nitrogen acquisition proceeds through organic-matter recycling rather than fixation

Given the relatively early successional state of the site, we screened the recovered MAGs and unbinned assemblies for nitrogen-cycling capacity. Nitrogenase potential was remarkably low: 43 *nifH* genes encoding nitrogenase were detected within the unbinned contigs, and only nine MAGs (5.7%) carried *nifH*, all of which were transcribed at near-background levels (**Figure 3C**). This scarcity aligns with reports of minimal diazotrophic capacity during early post-glacial succession [28,42]. It also supports the hypothesis that pioneer nitrogen demand in alpine forefields is met primarily by atmospheric deposition channeled through the rocky matrix, thereby alleviating the selective pressure for biological nitrogen fixation.

Consistent with an alternative acquisition route, many of our recovered MAGs encoded chitinases and ß-*N*-acetylhexosaminidases, equipping these microbes to degrade chitin and amino sugar polymers into glucosamine and *N*-acetylglucosamine. The subsequent metabolism of these monomers liberates free ammonia. At the level of genome-resolved potential, this corroborates the conclusion by Brankatschk *et al* (2011) that mineralization, specifically chitinolysis and proteolysis, rather than fixation, drives nitrogen turnover in initial forefield soils [43].

The distribution of other nitrogen-cycling genes further reflected reliance on recycling rather than *de novo* input. Markers for ammonia oxidation (*amoA*, n=4, 2.5%) (**Supp. Figure 5**) and nitrite oxidation (*nxrA*, n=4, 2.5%) were rare. In contrast, nitrite reductase (*nirK*) was detected in 19 MAGs (11.9%), representing nearly twice the prevalence of *nifH*, and nitrate reductases were similarly widespread (*narG*: n=8.8%, 10.7%; *napA*: n=13, 8.2%). These nitrogen-cycling pathways were carried by Actinomycetota (n=24), Pseudomonadota (n=10), Chloroflexota (n=6), Acidobacteriota (n=1) and Bacteriodota (n=1). Notably, transcripts for these genes were uniformly low across the community, indicating that nitrogen transformation operates as a background maintenance process at this successional stage relative to the dominant energy acquisition pathways driving trace-gas oxidation.

In contrast, carbon fixation potential was substantial. The *rbcL* gene encoding ribulose-1,5-bisphosphate carboxylase/oxygenase (RuBisCO), was the most widely distributed carbon-fixation marker screened, detected in 31 MAGs (19.5%), predominantly within Actinomycetota (23/31, 74.2%) and Chloroflexota (n=5, 16.1%), as well as 85 unbinned contigs. Notably, 83.9% of *rbcL*-carrying

MAGs (n=26/31) also encoded at least one trace-gas marker gene (Group 1 [NiFe]-hydrogenase, *coxL*). Both gene classes were transcriptionally active within these genomes, supporting a coupled metabolic mechanism wherein electrons derived from trace-gas oxidation power Calvin-Benson-Bassham (CBB)-mediated $CO_2$ fixation [44]. Additionally, a single Nitrospirota MAG (Palsa-1315, a comammox lineage) encoded a complete reverse tricarboxylic acid (rTCA) cycle, demonstrating that specialized nitrifiers occupy a complementary dark-fixation niche alongside the dominant CBB-utilizing generalists.

#### S3. Active MGEs carry mobilizable trace-gas machinery

Assemblies yielded 199 vOTUs and 138 pOTUs passing quality thresholds. Both were recovered mostly from the mesocosm experiment (vOTU 139/199; pOTU 116/138), which indicates an enrichment during the processing of sample for the mesocosm. Host-linked elements were assigned predominantly to Actinomycetota (pOTU=100; vOTU=36) and Pseudomonadota (pOTU=27; vOTU=28), reflecting the dominant community, with smaller assignments to Acidobacteriota (vOTUs n=4; pOTUs n=4), Bacillota (n=3); and Chloroflexota (pOTUs n=1). A large proportion of vOTUs (128/199) had no host prediction (**Supp. Figure 6A**). Twenty-eight pOTUs and 17 vOTUs were linked to MAGs recovered within this dataset.

Nearly all elements were transcriptionally active in the *in-situ* libraries (vOTU 160/199; pOTU=137/138), indicating that the extrachromosomal fraction of this community is not a dormant reservoir (**Supp. Figure 6B**). Expression was significantly correlated with the predicted host MAG for pOTUs (Spearman's $\rho = 0.38$ ($n = 28$, $p = 0.04$)), and vOTUs ($\rho = 0.72$ ($n = 17$, $p = 0.001$)) (**Supp. Figure 6C**), consistent with plasmid transcription tracking host physiology. Viral transcription also mirrored host transcriptional activity, indicating that the local virome operates predominantly through stable, temperate co-existence (lysogeny) or chronic, non-lethal infection rather than active host lysis, relying on the host consistent metabolism to maintain.

Three pOTUs carried central metabolic genes: a 150-kb contig assigned to *Variovorax* sp901827185 (*Burkholderiaceae,* Pseudomonadota) carrying heme-copper oxidases *CoxA* subunit 1, *CoxA* subunit 2 and *CyoA* involved in the respiratory transport chain, a 118-kb element from *Acetobacteraceae* (G192, Pseudomonadota) carrying two partial *CoxA* genes; and a 273.3-kb element from *Telluria beijingensis* (*Burkholderiaceae*, Pseudomonadota) carrying sulfide dehydrogenase (*fcc*). All elements/genes were actively expressed *in situ*, except *fcc* (**Supp. Figure 6D**).

##### S4. Acetylene inhibition successfully suppresses methanotrophy

In the first field campaign (2024; sites C and F), acetylene abolished net $CH_4$ uptake causing flux to decrease by 0.775 g $m^{-2}$ $yr^{-1}$ (C: -1.16; F: -0.389), from a mean uptake of 0.81 to 0.04 g $m^{-2}$ $yr^{-1}$ overall (**Supp. Figure 9**). Inhibition also altered the fluxes of both other gases. Net $H_2$ uptake decreased by 0.148 g $m^{-2}$ $yr^{-1}$ (C: -0.179; site F: -0.124) under treatment, consistent with direct inhibition of [NiFe] hydrogenases by acetylene. Net CO exchange shifted toward uptake by 0.190 g $m^{-2}$ $yr^{-1}$ (C: +0.163; F: +0.224), moving the sediment from a net CO source (-0.126 g $m^{-2}$ $yr^{-1}$) to a small net sink (+0.065 g $m^{-2}$ $yr^{-1}$), most notably at site F, where the flux changed from -0.165 to +0.059 g $m^{-2}$ $yr^{-1}$.
